# Glucagon and GLP-1 Accelerate Pseudo-Islet Assembly and Unmask Sex-Specific Islet Fragmentation Dynamics

**DOI:** 10.1101/2025.08.10.669547

**Authors:** Kaya Keutler, Stella Prady, Pamela Canaday, Craig Dorrell, Carsten Schultz

## Abstract

Pancreatic hormones are best known for their role in regulating blood sugar levels as well as islet cell function and proliferation. However, their impact on maintaining and inducing cell aggregation in culture remains under-explored. In this study, we investigated the effects of glucagon (GCG) and glucagon-like-peptide 1(GLP-1) on the formation and integrity of human islet clusters. Native human islets were dissociated and sorted into pure α-, β-, and δ-cell populations using antibody-based fluorescence-activated cell sorting (FACS). The sorted cells were then co-cultured with mouse endothelial MS1 cells in suspension to generate pseudo-islets of varying cell composition. Hormonal supplementation with GCG or GLP-1 versus blank was administered during the tissue culture phase. Hormone-treated pseudo-islets formed faster, dependent on the cellular composition and the sex of the donor. In parallel, we also exposed native islets, maintained in suspension without prior dissociation or sorting, to hormone supplementation. These islets exhibited accelerated fragmentation under hormone treatment compared to controls, again dependent on donor sex with islets from female donors fragmenting faster than from male donors. These findings suggest that GCG and GLP-1 enhance pseudo-islet formation and affect the structural integrity of native islets in a sex-specific manner, offering insights into islet biology and implications for diabetes research and therapy.

**Article Highlights:** We established a manipulatable, expandable human pseudo-islet platform to investigate islet morphogenesis, architecture, and intercellular signaling. We examined the contribution of individual α-, β-, and δ-cell populations and assessed how glucagon (GCG) and glucagon-like peptide-1 (GLP-1) modulate islet integrity in culture. In native islets, hormonal supplementation attenuated fragmentation in male donors but accelerated it in females. In pseudo-islets, cellular composition was the predominant determinant of maturation versus fragmentation, with donor sex exerting a secondary influence. We present methodological guidelines for generating and maintaining human pseudo-islets, thereby providing a framework to optimize donor selection, culture conditions, and experimental design in diabetes research.

## Introduction

Islets of Langerhans and their secreted hormones, glucagon, insulin, and glucagon-like-peptide 1 (GLP-1) are key regulators of glucose homeostasis. Dysregulation of hormone-secreting cells is the hallmark of diabetes. Therefore, it is essential to better understand how islet-secreted hormones contribute to islet well-being.

Apart from insulin, GLP-1 is among the most studied and has led to the approval of diabetes and weight control drugs such as Ozempic. GLP-1 works primarily through the GLP-1 receptor (GLP-1R, a G-protein coupled receptor), which, in the islet, is exclusively expressed by β-cells (1). GLP-1 has been shown to promote β-cell replication by inducing the expression of cell cycle regulators such as cyclin D1 (2). In addition to promoting β-cell proliferation, GLP-1 also exerts anti-apoptotic functions by activating survival signaling networks, including the PI3K/AKT pathway (3). GLP-1 increases β-cell mass in the islet, partially through β-cell proliferation, but also through inducing the conversion of α-cells into β-cells. This process involves the expression of fibroblast growth factor 21 (FGF21), facilitating the transdifferentiation process (4,5). Additionally, GLP-1 promotes insulin secretion by β-cells and suppresses glucagon secretion by α-cells through cAMP and the exchange protein activated by cAMP (EPAC) (6). Hence, local production of GLP-1 from pancreatic α-cells on demand may be beneficial to protect and regenerate β-cells. Healthy islets secrete more GLP-1 when stressed with high glucose (4). This has been interpreted as an adaptive mechanism and a way to boost insulin secretion from remaining β-cells (7).

In addition to GLP-1, α-cell-secreted glucagon (GCG) modulates β-cell function by regulating microtubule (MT) stability, impacting insulin secretion, and overall islet performance. In islets with a higher α/β-cell ratio, β-cells exhibit a less stable MT network that is associated with enhanced insulin release upon high glucose and/or depolarizing stimuli. This response is recapitulated by direct GCG stimulation (8–10). Mechanistically, GCG and GLP-1 activate cAMP production in β-cells, promoting both the nucleation of new MTs at the Golgi apparatus (thereby supporting insulin secretory granule biosynthesis) and the destabilization of existing MTs, which increases the pool of releasable insulin granules by potentiating Ca²⁺ influx and elevating granule sensitivity to Ca²⁺. Furthermore, β-cells in closer proximity to α-cells show more pronounced MT destabilization and insulin secretion, highlighting a spatial aspect of paracrine signaling that underlies the functional heterogeneity of islets (9–11). Further, as indicated by cell line co-culture to form pseudo-islets, α-cells secrete hormones that reduce oxidative stress and protect β-cell mass (12).

The importance of α-β-cell paracrine regulation is also indicated by the identification of GLP-1R nanodomains on the contact sites of α-cells and β-cells, where pre-internalization of GLP-1R at low glucose levels primes these β-cells for rapid and enhanced Ca²⁺ signaling and insulin secretion upon high-glucose stimulation. β-cells adjacent to α-cells exhibit earlier Ca²⁺ rises and nearly double the insulin release compared to those neighboring other β-cells (13).

GLP-1 and GCG are known to regulate islet size and architecture. Studies in transgenic mouse models have identified transcription factors such as Elk-1 and Egr-1, downstream of Ca²⁺ signaling, as key controllers of islet development and mass. Dysregulation of these pathways impairs islet architecture and glucose homeostasis. Therefore, both incretin and GCG signaling are critical for islet assembly during development and for preserving islet integrity in adulthood (14–16).

Human islet architecture is distinct from that of rodents, although it is suggested that within human islets, micro-domains follow the rodent islet organization of β-cells in the core and α-cells in the mantle (17–19). Little is known about the placement of δ-cells and the connection/placement of blood vessels. Although islets only represent 1–2 % of the pancreatic mass, they receive 6–20 % of the direct arterial blood flow to the pancreas (20). Based on measurements in the perfused rodent pancreas, Samols et al. observed a β → α → δ sequence, implying that δ-cells occupy the most downstream positions along the islet capillary network (21–23).

Over the past three decades, researchers have made substantial progress in characterizing the islet microenvironment and its signaling networks. However, these insights remain circumstantial because no model allows real-time study of islet organoid formation with manipulatable components. Without such a system, it is impossible to explain why native islets diverge in behavior from immortalized cell lines or to pinpoint which cell types or hormones drive those differences. Therefore, we have developed a pseudo-islet platform built from primary human cells that enables controlled assembly and manipulation of specific endocrine cell populations, providing a tool to dissect cell-type– and hormone-specific effects on islet development and function.

We demonstrate that immortalized pancreatic cell lines exhibit enhanced growth upon GCG or GLP-1 supplementation via their respective GPCRs. In contrast, native human islets showed no viability improvement and instead displayed accelerated fragmentation (24–26) in female, but not male, donor islets. To investigate these responses, we developed a controllable pseudo-islet model derived from human islets. These engineered pseudo-islets regained the hormone-induced growth benefit seen in cell lines, dependent predominantly on their cellular composition and less on donor sex.

## Methods

### Tissue culture of pancreatic cell lines

Mouse Insulinoma 6 (MIN6) cells were a kind gift from Dr. Miyazakis’s lab (Osaka, Japan). MIN6 cells were cultured in DMEM with 4.5 g/L glucose (Gibco, REF# 11965-092) supplemented with 70 µM β-mercaptoethanol (Gibco, REF# 21985-023) before use. Cells were passaged once per week and used within passages 17-30. αTC1 clone 9 cells were a kind gift from the lab of Dr. Grompe (Portland, OR, USA). αTC1 cells were cultured in DMEM with 1 g/L glucose (Gibco, REF# 11885084) that was supplemented with 15 mM HEPES (Cytivia, REF #AJ30727929), NEAA (Gibco, REF #11140-050), and 100 mg BSA (Sigma, REF# A7906). Cells were passaged every three days and used within passages 4-13.

### Monitoring cell growth in response to media supplementation

Pancreatic hormone-secreting cell lines (MIN6 (passages 17-30), and aTC1 cl. 9 (passages 4-13)) were used to monitor cell growth. Cells were kept in high-glucose DMEM (4.5 g/L glucose, Gibco, REF# 11965-092) and low-glucose DMEM (1 g/L glucose, Gibco, REF# 11885084), respectively. For supplementation, hormones were purchased from Sigma Aldrich and used at the concentrations described in Table 1.

**Table 1:** Overview of compounds and hormones used.

| **Media Supplementation** | **Final Concentration** |
| --- | --- |
| **GLP-1 (human)** | 0.1 µg/mL |
| **Glucagon (GCG, human, G3265)** | 0.1 µg/mL |
| **Somatostatin (SST)** | 100 nM |
| **Exendin 9-39 (GLP-1R antagonist, E7269)** | 0.1 µg/mL |
| **GLP-1R agonist (G8038)** | 0.1 µg/mL |
| **GCGRIIα** | 1 µM |

To generate a growth curve, cells were seeded in 12-well plates (Falcon, LOT #353043) at a density of 0.1 x 10^6^ cells/mL. Cell growth was determined by manually counting the cells in each well (triplicate per day) on T2-T6 post-seeding with a hemocytometer. Cells were diluted with trypan blue (REF #T8154). Additionally, cell counts were determined semi-automated using a Keyence EPI-microscope and FIJI software to analyze the generated images. For this imaging, on the respective days, cells were stained with 10 µg/mL Hoechst (REF#TG2611041) for 10 min, at 5 % CO_2_ and 37 °C. After washing, whole wells were imaged and analyzed using FIJI following the pipeline described in Supplemental Fig. 5.

Manually and semi-automatically, cell-count results were correlated by calculating the number of cells per area. For manually counted cells, the area corresponded to the number of cells counted for each square of the hemocytometer (1 mm^²^). Cells in an entire well (380 mm^2^) were counted semi-automatically. Manually counted cell numbers were compared to semi-automatically counted cells by multiplying by the dilution factor and then the area of the whole well.

### Handling of human pancreatic islets

Human pancreatic islets were provided by the NIDDK-funded Integrated Islet Distribution Program (IIDP) (RRID:SCR _014387) at City of Hope, NIH Grant # U24DK098085, and the JDRF-funded IIDP Islet Award Initiative (Study # BS562P & BS562). Additionally, human islets for research were provided by the Alberta Diabetes Institute IsletCore at the University of Alberta in Edmonton (http://www.bcell.org/adi-isletcore.html) with the assistance of the Human Organ Procurement and Exchange (HOPE) program, Trillium Gift of Life Network (TGLN), and other Canadian organ procurement organizations. Islet isolation was approved by the Human Research Ethics Board at the University of Alberta (Pro00013094) (27). All donors’ families gave informed consent for the use of pancreatic tissue in research. Islets from both distribution programs were shipped overnight and processed on the receiving day. If that was not possible, they were stored at 4 °C for no more than seven days. Native islets, when used as controls for pseudo-islets, were maintained in 1x CMRL 1066 media (Corning, REF# MT15110CV) supplemented with 10 % FBS, 10 mM HEPES (Cytivia, REF#AJ30727929), and 2 % L-glutamine (Gibco, REF#25030-081) with 1 % Pen/Strep (Gibco, REF# 15140122) and 1 µg/mL Amphoticerin B (Sigma, A2942, CAS # 1397-89-3) added just before use in culture. Human donor characteristics are listed in Supplementary Table S1.

### Tissue sources and pancreatic cell isolation

Human pancreatic islets from normal donors were obtained from the NIDDK-funded Integrated Islet Distribution Program (IIDP) at City of Hope (Study # BS562P & BS562). These were collected from approved, consented cadaveric organ donors from whom at least one other organ has been approved for transplantation and are exempt from human studies approval. Specimens were dispersed to single-cell suspensions by an 8–15 min digestion in 0.05% trypsin-EDTA (Gibco, REF#25-300-054) at 37 °C with dispersal by a p1000 micropipette every 3 min. The progress of the dispersion was checked by light microscopy. Undispersed tissue was removed with a 40-µm cell strainer (Fisherbrand, REF#22-363-547), and dissociated cells were stored on ice in holding buffer (CMRL1066 + 2% FBS + 0.1 mg/mL trypsin inhibitor (Sigma, REF#T9128) and 0.1 mg/mL DNAseI (Roche, REF#10104159001) before antibody labelling for FACS. The number of samples analyzed was primarily determined by material availability, but it was chosen to be sufficient for statistical analysis.

### FACS of dissociated human islets into pure α-, β-, and δ-cell populations

Dissociated cells were resuspended in holding buffer (CMRL1066 + 2% FBS + 0.1 mg/mL trypsin inhibitor (Sigma, REF#T9128) and 0.1 mg/mL DNAseI (Roche, REF#10104159001)) before the addition of antibodies. The antibodies used were: HIC1-2B4 A488 conjugated (Novus Biologicals, REF# NBP1-18946AF488) at a 1:100 dilution, HIC1-8G12 PE conjugated (provided by the Grompe lab) at 1:50 dilution, and anti-CD9 APC conjugated to APC (Thermo Fisher Invitrogen, MA1-10307) at a 1:20 dilution. Single antibody controls, A488 + PE, and all combined samples were incubated at 4°C for at least 20 min. After washing with cold CMRL1066, cells were resuspended in holding buffer, and 1 % propidium iodine (Sigma, REF#P4864) was added for live/dead distinction. Cell doublets were excluded by pulse width measurement, and propidium iodide staining was used to label dead cells for exclusion. Analysis was performed on a Cytopeia inFluxV-GS (Becton-Dickinson, BD) or a BD Symphony S6.

### Generation of pseudo-islets

Freshly sorted pancreatic cells were used to generate pseudo-islets. Pure cell populations were spun down directly after sorting and resuspended in 300 µL of media (CMRL1066 + 2% FBS + 1% Pen/strep, + 1 µg/mL Amphoticerin B). Based on the obtained cell numbers, concentrations in cells/µL were calculated and used to mix cells in specific ratios. If not indicated otherwise, cells were combined in a 5:1 ratio for β:α, 16:1 for β:δ, and 3:1 for α:δ. Additionally, pseudo-islet mixtures were combined with MS1 cells in a 1:10 ratio, with ten MS1 cells (kindly provided by the Grompe Lab, CRL-2279) for every one endocrine cell (unless otherwise specified). Cell mixtures were seeded in ultra-low attachment, flat-bottom 6-well plates (Corning, REF# CLS3471) in 3 mL of media (CMRL1066 + 2% FBS + 1 % Pen/strep, + 1 µg/mL Amphoticerin B). Cluster formation was allowed to happen for up to 14 days with hormone (0.1 µg/mL GCG or GLP-1) addition and culture media refill (100 µL/well to account for evaporation) every other day. Cluster formation was monitored by using a Keyence EPI-microscope in brightfield mode and FIJI software (see Supplemental Figure 5) to analyze the generated images.

### Supplementation of pancreatic cell lines, native islets, and pseudo-islets

For media supplementation, compounds listed in Table 1 were used in the concentrations described in column “Final Concentration”. Compounds were added every other day for the duration of the experiment. In case of native and pseudo-islets, 100 µL of media were refilled every other day to account for evaporation and to replenish nutrients, Pen/strep, and Amphoticerin B to prevent contamination.

### GSIS analysis

Samples (pseudo-islet mixtures and native donor islets) were removed from media and equilibrated by incubation for 1 h at 37°C, 5 % CO_2_ in KRHB + 2.8 mM glucose (pre-incubation). Cells were then transferred to fresh buffer (KRHB + 2.8 mM (basal) or 22.2 mM (stimulated)) glucose for 1 h at 37 °C, 5 % CO_2_. After incubation, buffers were collected and stored at 4 °C for subsequent ELISA analysis (Promega, Lumit, Insulin Kit REF#CS3037A05 and Glucagon Kit REF#W8020). Additionally, total human insulin and glucagon content was measured from an aliquot of RIPA-lysed (RIPA, ThermoFisher REF # 89900 + Protease Inhibitors, cOmplete Protease Inhibitor Cocktail tablets, Sigma, REF # 11697498001) clusters by human insulin/glucagon ELISA (Promega, Lumit). Statistical analyses of the results were performed using Graphpad Prism 10.5 (for analysis of variance tests) and Microsoft Excel (for mean and standard deviation).

### Immunofluorescent labeling of native and pseudo-islets

Native and pseudo-islets destined for immunohistochemistry were either fixed and stored or processed immediately for antibody staining. For storage, islets were fixed in 4 % paraformaldehyde (PFA) at RT for 15 min, then quenched with 30 mM glycine for 1 min. After centrifugation at 300 *× g* for 4 min, RT to pellet the islets, the SN was discarded, and the pellet was resuspended in 20 % (w/v) PEG 400 in PBS (storage solution) to fully cover the islets. These preparations were then stored at – 20 °C until use.

For direct staining, islets were fixed in 4 % PFA at RT for 1 h, pelleted at 300 *× g* for 4 min, and blocked in PBS containing 10 % FBS and 0.1 % Triton X-100 for 1 h. Following another 300 *× g* spin to remove blocking solution, islets were incubated without agitation in primary antibody mix—anti-insulin (ThermoFisher, REF#, 1:500), anti-GCGR (Abcam, REF#ab75240,1:250), and anti-GLP-1R (Iowa DSHB, REF#Mab 7F38,1:30)—in blocking buffer for 3–4 d at 4 °C. After a brief wash spin (300 *× g*, 4 min, RT), islets were incubated in secondary antibodies (Cy3-conjugated donkey anti-guinea pig (Jackson ImmunoResearch, REF#706-165-148), Alexa 488 goat anti-rabbit (Invitrogen, REF#A11034), and Alexa 647 goat anti-mouse (Invitrogen, REF#A21235), each at 1:1,000 in blocking buffer) at RT for 3 h or at 4 °C for up to 24 h. Finally, islets were spun once more, resuspended in ∼ 20 μL blocking buffer, and mounted on slides beneath a drop of DAPI-containing mounting medium (ThermoFisher, REF#P36966). Coverslips were applied and, once dry, slides were imaged on an Olympus F1200 confocal microscope using 405 nm, 488 nm, 559 nm, and 635 nm laser line excitation. Islets that were stored at – 20 °C were processed the same way for antibody staining, except for including additional washing steps beforehand to completely remove the storage solution.

### Software

For image processing, both FIJI (Supplemental Figure 5) and CellProfiler (Supplemental Figure 6) were used. To further streamline image analysis, FIJI macros were written for cell growth monitoring, native islet cluster counts, and pseudo-islet cluster counts. These scripts were deposited on GitHub and can be found under https://doi.org/10.5281/zenodo.16757638. CellProfiler pipelines were generated for each channel combination and are also available on GitHub (https://doi.org/10.5281/zenodo.16757674). For basic calculations, such as mean and standard deviation, Microsoft Excel was used. For complex statistical evaluation, GraphPad Prism 10.5 was used.

### Statistical Analysis and Reproducibility

Donors were stratified by sex (female vs. male) to account for sex-specific differences; no other donor characteristics (*e.g.*, BMI, age, A1C) were used for grouping. For native islets, each condition was assayed in two technical replicates unless otherwise noted. Pseudo-islet experiments followed the same scheme with two technical replicates per condition whenever sort yields permitted. In total, experimental results from 10 female and 14 male donors served as biological replicates. All statistical comparisons were made by two-way ANOVA with Benjamini–Hochberg correction for multiple testing.

## Results

### GCG and GLP-1 promote the growth of pancreatic cell lines

MIN6 (β-cell) and αTC1 clone 9 (α-cell) lines were maintained in medium supplemented with 0.1 µg/mL GCG and/or GLP-1. Both peptides significantly increased cell growth compared with controls, as assessed by increases in Trypan Blue exclusion and Hoechst-stained cell counts. For quantification, cells at each condition (CTRL, + GCG, + GLP-1) were seeded into a 12-well cell culture plate, and cell numbers were assessed on days 2, 4, and 6 post-seeding (see Fig. 1a). The observed increase in cellular growth was consistent with current literature as summarized by Zheng *et al.* (28). However, the magnitude of the response depended on the cell type: GCG elicited the greatest proliferation in MIN6 cells, whereas GLP-1 drove the strongest increase in αTC1 clone 9 cells. To dissect receptor-mediated mechanisms, we treated cells with selective agonists and antagonists for the GCGR and GLP-1R receptors. Antagonism of GCGR with the small molecule GCGRIIα (a selective, non-competitive, high-affinity GCGR antagonist) abolished the proliferative effect observed with GCG, without deviating from control conditions when given alone. Further, inhibition of GLP-1R by Exendin 9-39 blocked GLP–1–induced α-cell growth, whereas Exendin 9-39 alone had no impact on basal proliferation of αTC1 clone 9 cells (Fig. 1).

**Figure 1:**
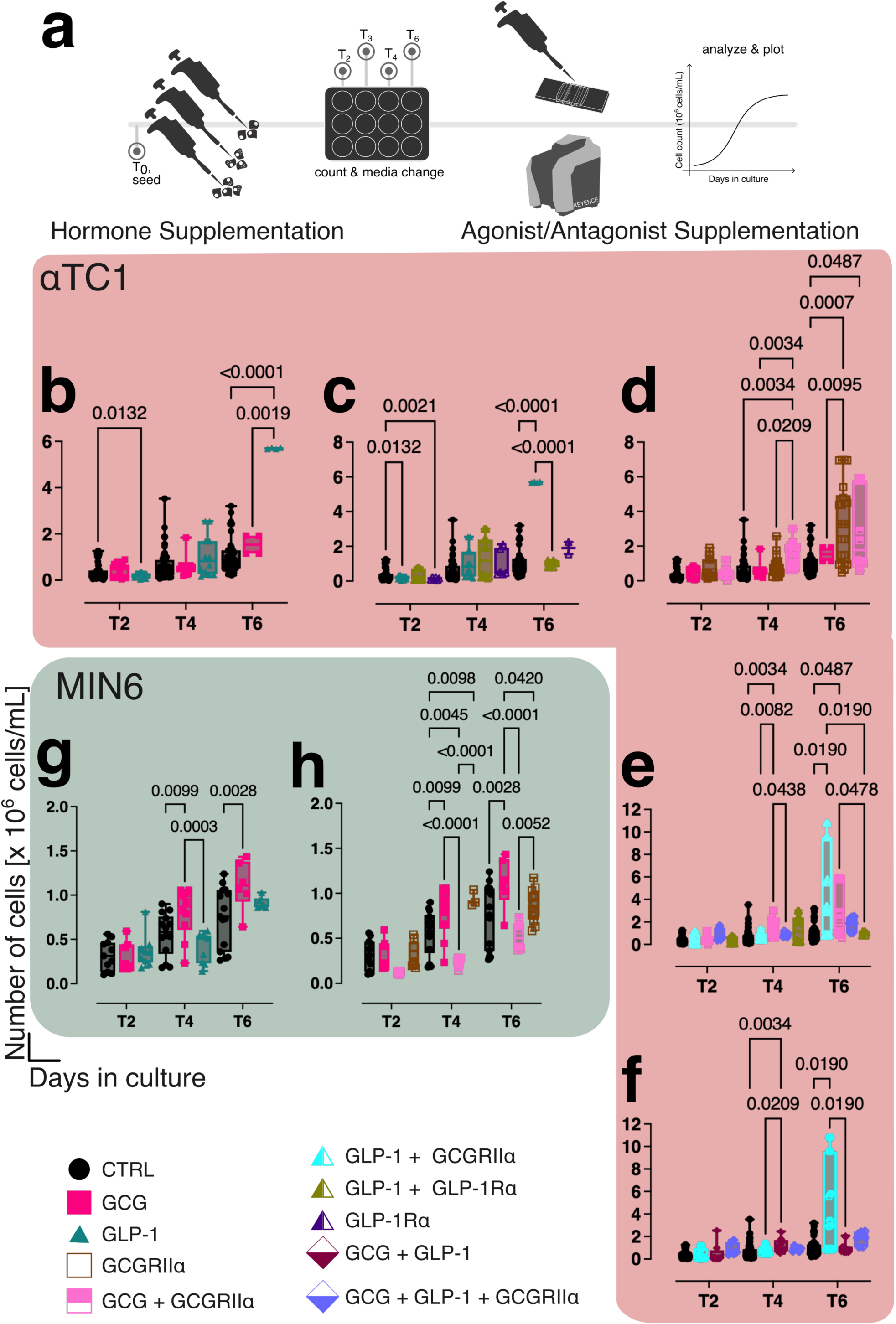
Hormone supplementation influences the Growth of pancreatic cell lines. a) Pancreatic cell lines, MIN6 representing β-cells and αTC1 representing α-cells, were seeded in 12-well plates and supplemented with single pancreatic hormones or combinations thereof. Growth monitoring was carried out for up to 6 days post-seeding with media exchange, hormone addition, and cell counts every day starting from T2 post-seeding (T3 and T5 counts are not shown for a better overview). b-f) αTC1 cells (red panel) supplemented with 0.1 µg/mL of hormones (GCG (pink) or GLP-1(teal)), 0.1 µg/mL of GLP-1Ragx (GLP-1R antagonist, purple triangle), 1 µM GCGRIIα (GCGR antagonist, brown) or combinations thereof (see legend for color associations). g & h) MIN6 cells (green panel) supplemented with 0.1 µg/mL of hormones (GCG (pink) or GLP-1(teal)), 1 µM GCGRIIα (GCGR antagonist, brown) or combinations thereof (see legend for color associations). Black represents CTRL conditions, which is the respective cell type in regular media without any additional supplementations. Statistical analysis was done by two-way ANOVA or mixed model (data dependent) for matched values in subcolumns. The Benjamini-Hochberg correction was used to adjust for multiple comparisons. p values < 0.05 (considered the cutoff for statistical significance) are listed on the graphs.

### GCG and GLP-1 sex-dependently alter the fragmentation of native human islets in culture

We next repeated the cell culture experiments using native human islets. Islet cells did not proliferate without or in response to 0.1 µg/mL GCG or GLP-1, consistent with the reported minimal replicative capacity of adult β-cells ex vivo (29). However, isolated primary islets are known to fragment over time in standard culture (as exemplary indicated in Fig. 2a), with a rapid shift toward smaller islet particles occurring within the first 24 h (24–26). Fragmentation proceeded even faster when we maintained islets in culture for up to 10 d with cluster counts at 2-3, 6-7, and 9-10 d post-seeding (Fig. 2b (female) and c (male)). When we supplemented native human islets with GCG or GLP-1, this fragmentation was significantly accelerated compared with untreated controls. This was reflected by an increase in 30 µm diameter cluster counts from 2 to 10 d post-seeding, with percent increases of 87.1 % in females and 130.9 % in males under control conditions, 68.2 % in females and 85.8 % in males with GCG, and 137.2 % in females and 99.8 % in males with GLP-1 (Fig. 2d (male) and Fig. 2g (female)). Additionally, changes in the proportion of subclusters (<30 µm diameter; Fig. 2e (male) and Fig. 2h (female)), expressed as a percentage of control, further supported ongoing fragmentation. In males, subcluster abundance in the + GCG condition was elevated at 2 d (114.2 %) but declined by 6 and 10 d (77.5 % and 86.3 %, respectively). A similar pattern was observed with + GLP-1 (117.8 %, 91.2 %, and 83.6 %). In contrast, female islets showed consistently elevated subcluster proportions. With + GCG, values were 192 % at 2 d, 129 % at 6 d, and 147.6 % at 10 d. With + GLP-1, subclusters remained high across all time points (184.7 %, 199.4 %, and 193 %). Both cluster and subcluster counts were assessed by the analysis of EPI microscopy-captured images with a FIJI pipeline described in Supplemental Figure 4. Therefore, hormone-treated islets exhibited an exaggerated loss of structural integrity during long-term culture when treated with α-cell-derived hormones.

**Figure 2:**
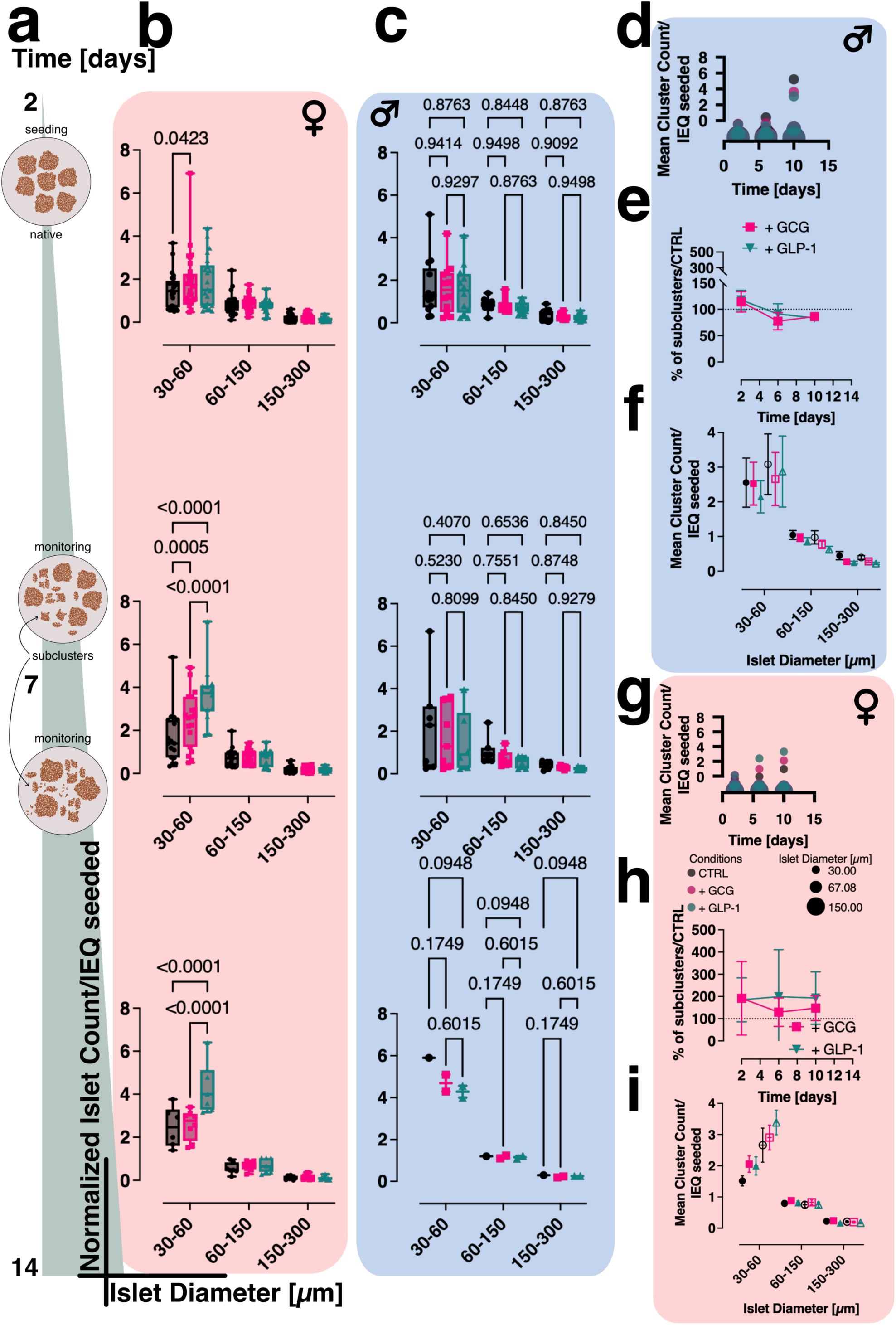
Response to hormone supplementation of native islets is influenced by donor sex. Growth and formation of native islets were monitored by light microscopy, monitoring islet diameter [µm] for up to 10 days post-seeding. a) Experimental timeline and modeling of expected behavior in culture. b) Top row corresponds to 2-3 days post-seeding, middle to 6-7 days, and bottom row to 9-10 days. Data was pooled by donor gender (n = 35 replicates for female donors (b) and n = 16 replicates for male donors (c)). Hormone supplementation with 0.1 µg/mL of either GCG (pink) or GLP-1 (teal) was compared to the CTRL condition in black. d) and g) mean cluster counts for cluster sizes of 30 µm – 150 µm normalized to the number of islet equivalents (IEQ) seeded, plotted over time in culture for male (d) and female (g) donors. Spheres are colored by condition (black = CTRL, pink = + GCG, and teal = + GLP-1), and the size corresponds to the size in islet diameter (µm). e) and h) Subclusters, defined as clusters < 30 µm, expressed as % of control condition and plotted over time in culture for + GCG (pink) and + GLP-1 (teal) for male (e) and female (h) donors. f) and i) mean cluster count normalized per IEQ seeded for islet diameter ranging from 30 – 150 µm. Filled symbols for CTRL (black), + GCG (pink) and + GLP-1 (teal) correspond to clusters counted 2-3 days post-seeding. Clear symbols for CTRL (black border), + GCG (pink border), and + GLP-1 (teal border) correspond to clusters counted 6-7 days post-seeding. Statistical analysis was done by two-way ANOVA with a Benjamini-Hochberg correction for multiple comparisons. All p-values are indicated on graphs, space-permitting, otherwise only significant p-values are shown; p < 0.05 was selected as being statistically significant.

The effect of hormone supplementation on islet fragmentation was donor sex–dependent. In female islets, supplementation with GCG and GLP-1 led to increases in 30 µm diameter cluster counts to 78.3 % and 157.5 % of control values, respectively (Fig. 2g and i). In contrast, male islets showed more modest responses, with values reaching 65.6 % (+ GCG) and 76.2 % (+ GLP-1) of control (Fig. 2d and f). A similar trend was observed in the proportion of subclusters, where female islets exhibited elevated fragmentation under both treatments (μ = 158.3% with GCG and μ = 186.5% with GLP-1, relative to control; Fig. 2h). In comparison, male islets fragmented over time but remained at or below control levels, with subcluster proportions of μ = 88.8 % (+ GCG) and μ = 92.3 % (+ GLP-1) (Fig. 2e). All other donor-to-donor differences, such as age, ethnicity, BMI, and A1C, were ignored, and donors were only pooled based on their sex. Islets from male donors maintained a fragmentation profile like untreated control islets or showed improvement despite the presence of GCG or GLP-1, whereas female islets exhibited markedly increased fragmentation in response to both treatments. Therefore, our data indicate that GCG and GLP-1 promote structural destabilization of human islet clusters in a sex-dependent manner.

### Primary human islet cells form pseudo-islets and respond to pancreatic hormone supplementation

Native human islets did not exhibit growth or any other beneficial effect in response to GCG or GLP-1. However, they represent fully differentiated tissue and do not mimic the dynamic process of islet formation. In contrast, cell lines that form clusters during proliferation are considered non-physiological. To address this limitation, we developed a pseudo-islet model that recapitulates the early assembly of islets from primary human α-, β-, and δ-cells. This model not only enables the study of cell-cell interactions and hormone effects during islet reconstitution but also provides a standardized, human cell–based platform for functional studies.

We generated pseudo-islets after dissociating native human islets into single cells and sorting them into pure α-, β-, and δ-cell fractions by fluorescence-activated cell sorting (FACS) using a pre-established procedure from the Grompe lab (30) (Fig. 3a, Supplemental Fig. 2).

**Figure 3:**
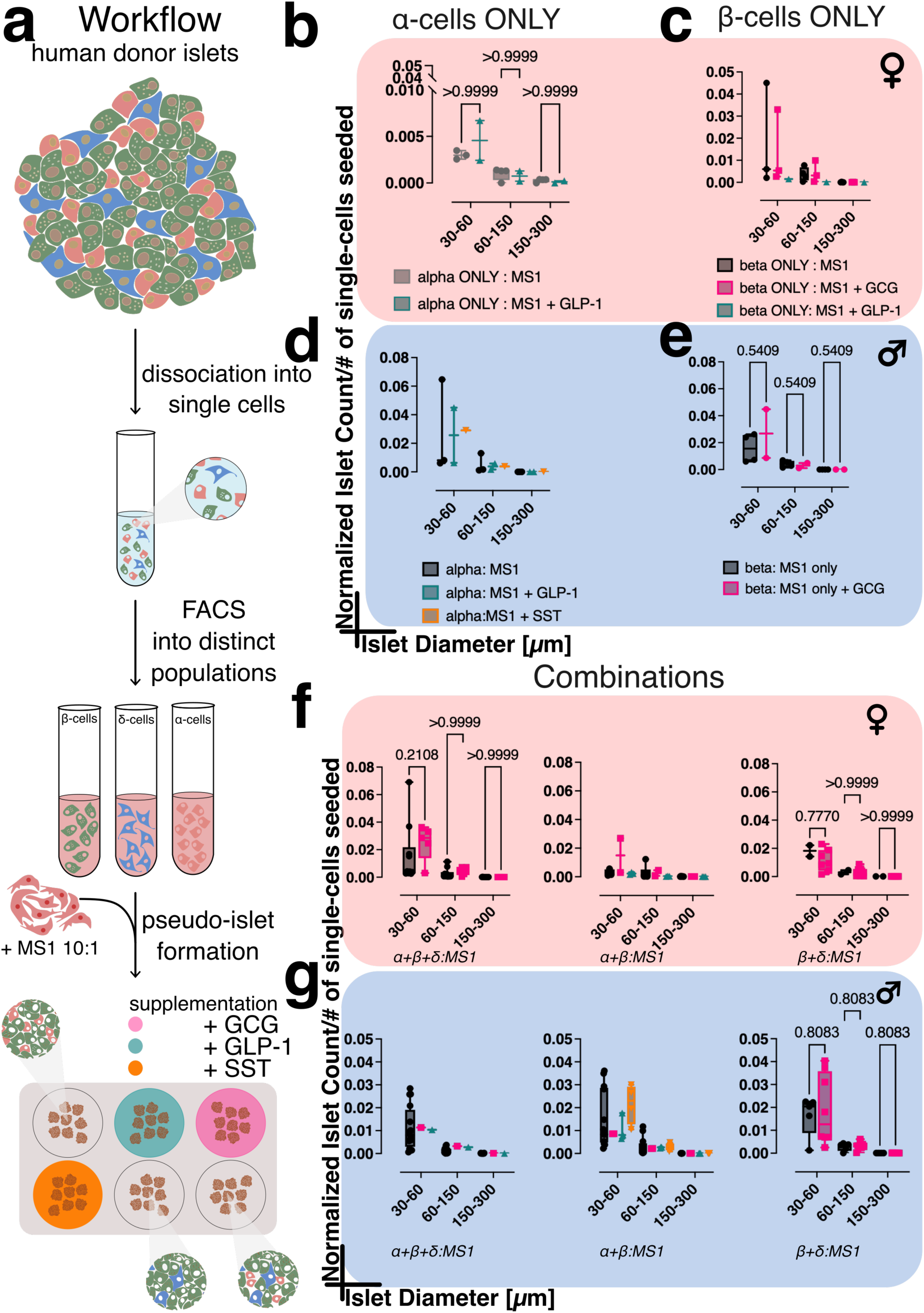
Human-donor-derived α-cells, β-cells, and combinations thereof form cell clusters (pseudo-islets) that are responsive to pancreatic hormone supplementation. a) Workflow: Human donor islets derived from both male and female donors were dissociated into single cells, stained with HIC1-2B4 AF488 (pan-endocrine marker), HIC1-8G12 PE (non-β-endocrine cell marker), and CD9 APC (predominantly δ-cell marker), and sorted into distinct α-, β-, and δ-cell populations via FACS. These populations were then used to form pseudo-islets either from single populations (b) and d) for α-cells ONLY and c) and e) for β-cells ONLY) or combinations thereof (f and g). Pseudo-islets were formed in ultra-low attachment 6-well plates and combined in a 1:10 ratio of endocrine to Mile Sven 1 (MS1) cells (mouse pancreatic endothelial cells). Hormone supplementation consisted of 0.1 µg/mL GCG (pink), 0.1 µg/mL GLP-1 (teal), and 100 nM somatostatin (SST, orange), compared to the control (CTRL, black), with supplements administered every other day. Growth and formation of pseudo-islets depicted in all subpanels are from 2-3 days post-seeding via EPI-light microscopy. Donors were pooled by sex; panels b), c), and f) correspond to female donors and panels d), e), and g) to male donors. Statistical analysis was done by two-way ANOVA with a Benjamini-Hochberg correction for multiple comparisons. P-values are indicated for each comparison; p < 0.05 was considered the cutoff for statistical significance.

To support three-dimensional re-aggregation, we mixed defined combinations of endocrine cells with MS1 pancreatic endothelial cells at a 1:10 endocrine:non-endocrine ratio, a method known to promote free-floating pseudo-islets with enhanced structure and function in ultra-low attachment plates (32) (Fig. 3a). We anticipated that pseudo-islets would form over time from clustering of single cells, beginning as small subclusters (< 30 µm) and growing into larger clusters within defined size bins (30–60, 60–150, or 150–300 µm) (Fig. 4a). However, pseudo-islet behavior in culture was less intuitive to interpret compared to native islets due to differences in normalization. Native islet cluster counts were normalized to IEQs seeded per well, with values <1 indicating loss or fusion, >1 indicating fragmentation, and interpretation further refined by subcluster percentage. In contrast, pseudo-islet cluster counts were normalized to the number of single cells seeded. Here, changes in cluster counts and subcluster percentages reflected fragmentation, fusion, or maturation. A summary of interpretative scenarios is provided in Supplemental Table 2.

**Figure 4:**
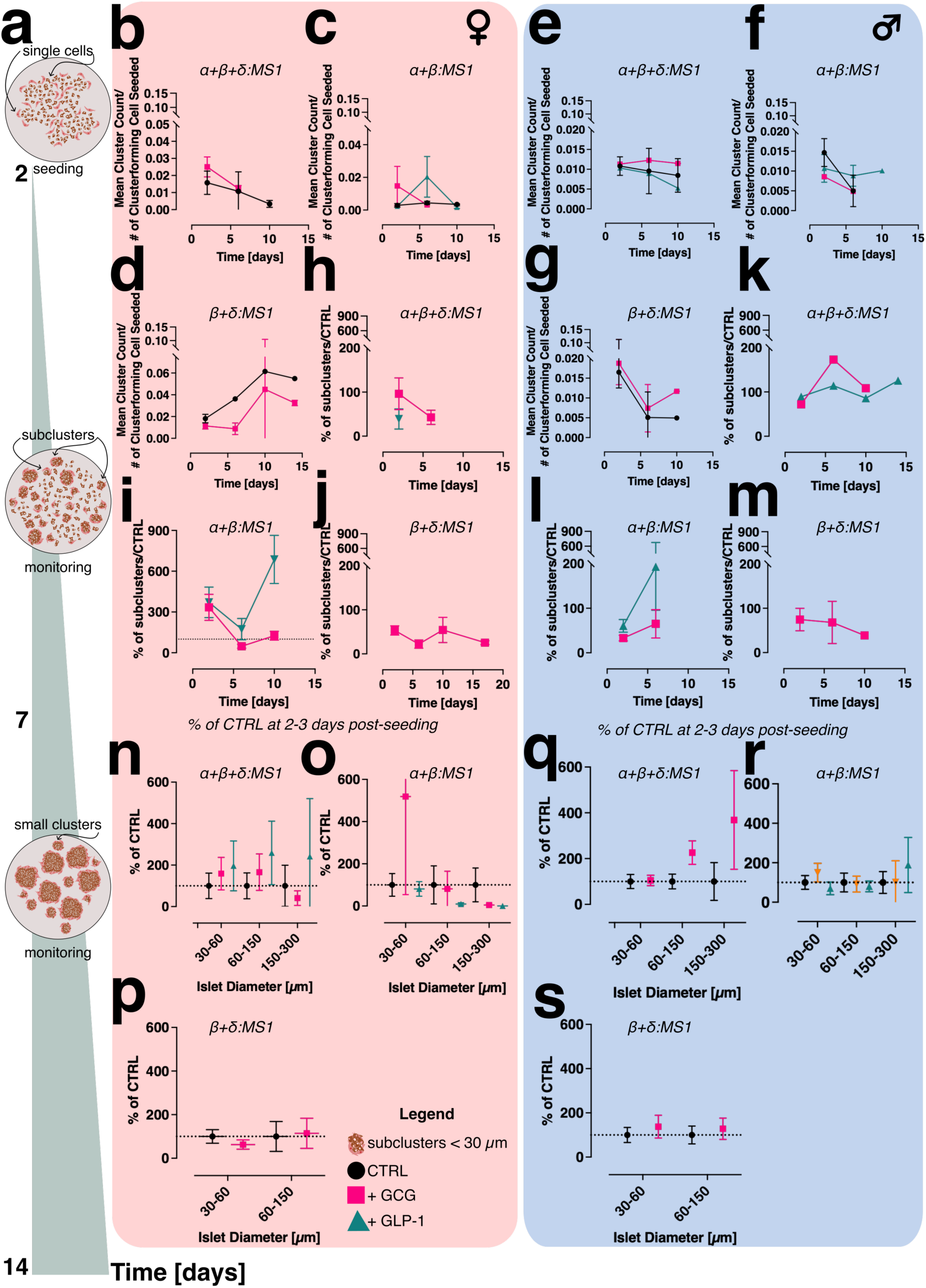
Pseudo-Islet Formation is influenced by cell composition, hormone supplementation and donor sex. a) theory, depicting the changes occurring to pseudo-islets as they mature in culture over time. b), c), and d) female “dynamic growth” curves. e), f), g) Male “dynamic growth” curves. Both are expressed as mean cluster count/seeded cluster-forming cell number. h-j (Female) and k-m (male)) % of subclusters normalized to control for pseudo-islet mixtures as depicted. n-p (Female) and q-s (male)) pseudo-islet growth at 2-3 days post-seeding expressed as % CTRL. Statistical analysis was done by two-way ANOVA with a Benjamini-Hochberg correction for multiple comparisons. P-values are indicated for each comparison; p < 0.05 was considered the cutoff for statistical significance. Error bars represent the SEM.

Pseudo-islets were found to form robust, compact spheroids as observed by brightfield microscopy (Supplemental Fig. 7). Those containing β-cells displayed the most islet-like morphology, characterized by a spheroid structure with well-rounded edges and an amber hue under brightfield microscopy. However, pseudo-islets also formed from pure β- or α-cells (Fig. 3b–e) or their combinations (Fig. 3f–g). Supplementation with GCG or GLP-1 altered pseudo-islet formation and growth compared to unsupplemented controls. These effects depended on cellular composition, donor sex, and the type of supplementation. In female-derived β+δ pseudo-islets, cluster size shifted progressively toward smaller aggregates under both conditions, with complete loss of 150–300 µm clusters by day 10 (+ GCG) or day 14 (control) (see Supplemental Fig. 3). Subcluster percentages remained below 100 % under + GCG (Fig. 4j), suggesting maintenance of intermediate-sized clusters. In contrast, male-derived β+δ pseudo-islets retained larger clusters (60–150 and 150–300 µm) over time, with minimal fragmentation; subcluster percentages remained lower under + GCG than control (Fig. 4m). To complement size bin analyses, dynamic growth curves (30–300 µm) were used. In female pseudo-islets, + GCG suppressed growth by up to 120 % compared to control (Fig. 4d). In males, both conditions showed an early decline, but + GCG later promoted recovery (62.2 % by day 10; Fig. 4g). Full interpretations for other combinations are summarized in Supplementary Table 2.

Like native islets, pseudo-islets show sex-specific differences, although more nuanced. Our results indicate that both female- and male-derived pseudo-islets respond to GCG and GLP-1 supplementation during early stages of pseudo-islet formation, with evidence of increased aggregation or maturation depending on the condition. However, over longer culture periods (9–10 d), signs of fragmentation emerge, particularly in female-derived pseudo-islets treated with GLP-1 (Fig. 4i). Increased subcluster percentages and a decline in cluster counts indicate this. In male-derived pseudo-islets, fragmentation tends to occur earlier, often by 6–7 d, and is especially pronounced in the + GLP-1 condition (Fig. 4e and f). These findings suggest that the structural trajectory of pseudo-islets is cell composition-dependent, sex-dependent, and hormone-specific, with early hormone-induced stabilization in some combinations giving way to later-stage fragmentation.

### Sex-specific analysis of islet hormone secretion and receptor expression in native and pseudo-islets

Cultured native and pseudo-islets were assessed following the workflow outlined in Fig. 5a. For native islets, fragmentation was monitored, while for pseudo-islets, formation was observed. Brightfield images were acquired every 2–3 days using EPI light microscopy. After 7 days, intact native islets and formed pseudo-islets were manually harvested for functional assays of hormone secretion. Insulin and glucagon secretion were quantified using the Promega Lumit detection kits (Supplemental Fig. 4). After functional assessment, remaining islets were fixed in PFA and processed for immunofluorescent staining to detect glucagon receptor (GCGR) and glucagon-like peptide-1 receptor (GLP-1R) expression. Antibodies were used at concentrations previously validated by the Hodson laboratory (13), and the staining protocol was generously provided by Dr. M. Golson (private communication).

**Figure 5:**
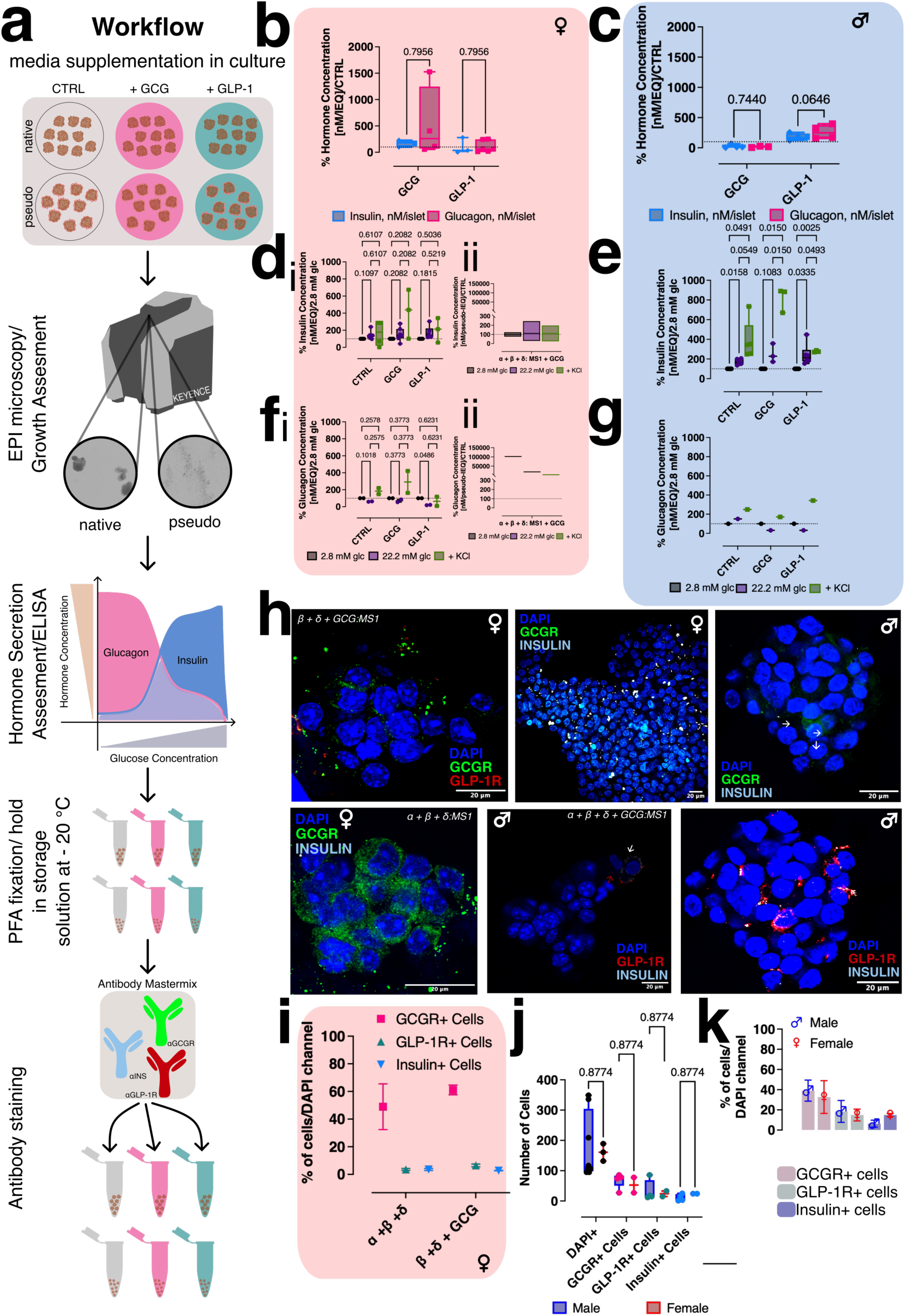
Sex-specific functional responses and receptor expression in native and pseudo-islets. a) Workflow schematic: steps taken to functionally characterize native and pseudo-islets for hormone secretion and receptor expression, validated by immunohistochemistry. b and c) total hormone content for glucagon (pink) and insulin (blue) for native islets from female and male donors, respectively. d_i_ and e) mean insulin concentration assayed from native islet supernatant in response to 2.8 mM (black), 22.2 mM glucose (purple), and 30 mM KCl (green) for female and male donor islets, respectively. fi and g) mean glucagon concentration assayed from native islet supernatant in response to 2.8 mM (black), 22.2 mM glucose (purple), and 30 mM KCl (green) for female and male donor islets, respectively. d_ii_ and f_ii_) mean insulin and glucagon concentration, respectively, measured from the supernatant of pseudo-islets formed from α+β+δ-cells (+/- GCG) in response to 2.8 mM glucose (black), 22.2 mM glucose (purple), and 30 mM KCl (green) for female donors only. For all ELISA results, values are normalized per IEQ used in the experiment and expressed as % of response to 2.8 mM glucose or CTRL. h) Representative immunofluorescent images for both native and pseudo-islets from female and male donors stained with DAPI (blue), GCGR (neon green), GLP-1R (dark red), and insulin (white). i) Percentage of GCGR+, GLP-1R+, and insulin+ cells normalized to DAPI+ (=total cells) for pseudo-islets derived from female donors. j) cell count comparison between native islets from female (red) and male (blue) donors. k) cell counts from j) expressed as % of DAPI+ cells for both female (red) and male (blue) donors. Statistical analysis was done by two-way ANOVA with a Benjamini-Hochberg correction for multiple comparisons. P-values are indicated for each comparison; p < 0.05 was considered the cutoff for statistical significance. Error bars represent the SEM.

To identify β-cells, co-staining for insulin and GLP-1R was performed following the current understanding in the literature that GLP-1R expression is restricted to insulin-producing β-cells (1). This approach doubled as a validation of the observed staining, as INSULIN+ cells were expected to overlap with GLP-1R+ cells.

Native islets from both female (n=9, Fig. 5b) and male (n=3, Fig. 5c) donors showed no significant differences in total insulin or glucagon content under culture conditions supplemented with either glucagon (+ GCG) or GLP-1 (+ GLP-1), suggesting that exogenous hormone supplementation does not alter hormone synthesis in native islets. Hormone secretion was normalized to the number of IEQ and measured at basal glucose (2.8 mM), high glucose (22.2 mM), and in the presence of 30 mM KCl to assess maximal depolarization-induced release (stimulation protocol detailed in Supplemental Fig.4). Basal glucose-induced secretion was used as a baseline to express % of hormone secretion in response to high glucose and 30 mM KCl. Islets from female donors exhibited no significant difference in insulin (Fig. 5d_i_) or glucagon (Fig. 5f_i_) secretion across these conditions. In contrast, male donor islets showed a canonical secretory pattern, with stepwise increases in insulin release in response to increasing glucose concentrations, and a peak following KCl stimulation (Fig. 5e). Aside from the CTRL condition, a similar canonical pattern was observed for glucagon secretion. Here, decreasing values are expected with increasing glucose concentration. The + KCl condition is still expected to reflect total hormone release. However, not enough data could be collected from male donors to generate a strong data foundation. For pseudo-islets, data availability was even more scarce, considering the elaborate path of their generation. Despite that, some female donor-derived pseudo-islets could be analyzed for both insulin and glucagon secretion (Fig. 5d_ii_ and f_ii_, respectively). Here, secretion levels were normalized per pseudo-IEQ used in the experiment and expressed as % secretion of the control (= α+β+δ:MS1). Again, a canonical pattern represents increased insulin release with increasing glucose concentration but decreased glucagon release under the same conditions.

Receptor expression: To better understand the differences between islets from male and female donors, we quantified receptor expression levels by immunohistochemistry (Fig. 5a). Native and pseudo-islets were immunostained and imaged by confocal microscopy using single-plane and z-stack acquisition. Representative slices from z-stack projections are shown in Fig. 5h. Image processing and quantitative analysis were conducted using Fiji and a custom pipeline implemented in CellProfiler (see Supplementary Fig. 6). Nuclei were identified by DAPI staining and expanded by 15 pixels to approximate cytoplasmic boundaries. Fluorescent signals for GCGR, GLP-1R, and insulin were mapped onto DAPI-positive regions, and only cells with overlapping signals in the cytoplasmic region were classified as positive for each marker. Cell counts were quantified for each condition and normalized to the total number of DAPI+ nuclei. Both native and pseudo-islets from male and female donors exhibited similar cellular GPCR composition, with approximately 40 % GCGR+ cells, 20 % GLP-1R+ cells, and 15 % INSULIN+ cells (Fig. 5j, Fig. 5k). Notably, pseudo-islets displayed more intense immunostaining than native islets, presumably due to improved antigen accessibility (Fig. 5i).

## Summary

Collectively, our findings reveal several novel insights into islet biology. First, in MIN6 and αTC1 cell line experiments, GCG selectively enhanced β-cell (MIN6) proliferation without affecting α-cell (αTC1) growth, underscoring the distinct responses of different islet cell types. By using selective antagonists for the glucagon receptor (GCGR) and GLP-1 receptor (GLP-1R), we confirmed that these effects are mediated directly through their respective receptors, thereby establishing a mechanistic basis for hormone-driven islet cell expansion.

When translated to human islets, hormone supplementation uncovered a sex dependence in the structural stability of native islets, while in reconstituted pseudo-islets, the growth response to GCG and GLP-1 was less dependent on the donor sex but varied with time and according to the cellular composition of the pseudo-islet.

After evaluating islet preparations from multiple donors and isolation centers, we established the following best practices to generate healthy pseudo-islets. Native islets should have minimal culture time before shipment (each islet offer specifies duration) and must not be precultured before FACS. Reserve a subset of fresh islets as native controls before dissociation to account for heterogeneity. Process incoming islets on the day of receipt whenever possible; otherwise, store them at 4 °C and use them within seven days. Dissociate using 0.05% trypsin-EDTA (other digestion methods may harm antibody epitopes), monitoring under light microscopy to stop digestion once only single cells remain and no large clusters persist. After sorting, immediately resuspend cells in fresh media containing penicillin/streptomycin and amphotericin B to prevent contamination. Standard cell culture plates promote adhesion; therefore, seed cells in ultra-low attachment plates together with MS1 endothelial cells at a 1:10 endocrine-to-MS1 ratio. MS1 cells support 3D reaggregation but have been reported not to proliferate in suspension. Clusters begin forming within 2–3 days and continue maturing up to 14 days. Depending on the cellular composition of the pseudo-islet. Additionally, most combinations show the complete spectrum of islet diameters measured (30-300 µm) at the first 2-3 days post-seeding timepoint, suggesting that pseudo-islet formation occurs quickly. Therefore, we would suggest including an earlier observation time point. Sometimes, for example, for both male and female β+δ combinations, longer culture yields larger pseudo-islets but may increase functional stress.

Hormone supplementation effects were sex-dependent. Female donor–derived native islets fragmented in response to hormone supplementation and should not be cultured with supplements. Male donor–derived native islets, in contrast, benefited from supplementation, which slowed fragmentation relative to controls. For pseudo-islets, β+δ combinations showed the most consistent cluster formation. In female donor–derived pseudo-islets, β+δ clusters remained stable for up to 10 days without significant fragmentation. Glucagon supplementation was tolerated, but unsupplemented conditions yielded better outcomes. In male donor–derived β+δ pseudo-islets, early time points were optimal, with robust cluster formation within the first 5 days. Glucagon supplementation improved stability and is preferred over unsupplemented conditions.

Certain combinations were more prone to fragmentation and should be avoided or time-limited. In male donor– derived α+β+δ pseudo-islets, GLP-1 supplementation promoted fragmentation, whereas control and GCG-treated conditions remained stable up to 10 days, with GCG being preferred. Male donor–derived α+β pseudo-islets fragmented under all conditions and should be used, if at all, within the first 3 days of culture. In contrast, female donor–derived α+β pseudo-islets formed clusters with both GCG and GLP-1 supplementation, with GLP-1 yielding more mature aggregates. However, culture duration should not exceed 7 days, as fragmentation becomes evident by days 9–10.

## Discussion

We demonstrate that supplementation with GCG and GLP-1 differentially affects growth, native islet fragmentation in culture, and pseudo-islet formation, with the magnitude of the effect depending on the specific cell type(s) present. In pancreatic cell lines, GLP-1 supplementation was growth beneficial for both α- and β-cells, whereas GCG supplementation appeared to selectively support β-cells (Fig. 1). This cell type-specificity is likely explained by the endogenous secretion of GCG by α-cells, which may saturate GCGR signaling in these cells, rendering additional GCG ineffective. Conversely, β-cells that do not have endogenous GCG remain sensitive to exogenous GCG.

Supporting this interpretation, the beneficial effects of GCG on β-cells were abolished when GCGR was pharmacologically inhibited using GCGRIIα. This highlights that the observed improvements in β-cell growth following GCG supplementation are dependent on intact GCGR signaling.

In αTC1 cells, GCG alone (at 0.1 µg/mL; ≈30 nM) produces only a modest proliferative effect, compared to GLP-1 at the same concentration. When both hormones are combined, GCG competes for GLP-1R binding (EC₅₀ for GLP-1R–mediated cAMP production is ∼36 nM (31)), thereby displacing GLP-1 and acting as a partial agonist. This likely suppresses the robust growth otherwise induced by GLP-1. When the GCGR antagonist GCGRIIα is added at 1 µM, GCGR signaling is fully blocked, diverting both exogenous GCG and any endogenous ligand to GLP-1R, and this diversion elicits a strong proliferative response even in the absence of added GLP-1. Therefore, in the presence of both GCG (∼ 30 nM) and 1 µM GCGRIIα, GCG cannot signal through GCGR and instead engages GLP-1R. However, because GCGs intrinsic efficacy at GLP-1R is lower than that of GLP-1, it slightly blunts the maximal proliferative signal seen with GCGRIIα alone yet still drives substantially greater growth than GCG treatment by itself.

Interestingly, sex-specific differences emerged in response to both islet culture and hormone supplementation. Fragmentation rates differed between male and female donors. Male native islets benefited from both GCG and GLP-1 supplementation, as it slowed fragmentation compared to control conditions. However, female-derived native islets seemed to fragment faster under supplemented conditions, with + GLP-1 being the most fragmentation-prone condition. Female native islets being more sensitive to hormone supplementation and responding with faster fragmentation could represent a physiological adaptation unique to female islets. Estrogen and other sex hormones have been shown to enhance β-cell survival, proliferation, and insulin secretion, particularly under stress conditions (32,33). Across their lifespan, females experience substantial hormonal fluctuations during the menstrual cycle, pregnancy, and menopause. The capacity for dynamic reorganization of islet architecture, including fast aggregation and reaggregation, may therefore serve a protective or regulatory function in response to these hormonal shifts. While the precise mechanistic link between hormone-driven plasticity and islet dynamics remains to be defined, these findings underscore the importance of considering sex as a biological variable in islet physiology and highlight the potential for sex-specific therapeutic strategies in diabetes.

Pseudo-islet in culture behavior was less sex-specific and driven more by the cellular makeup of the pseudo-islets, although male-donor-derived and female-donor-derived pseudo-islets did show differences.

Both female- and male-derived pseudo-islets respond to GCG and GLP-1 supplementation during early stages of pseudo-islet formation, with evidence of increased fragmentation or maturation depending on the condition. Pseudo-islets derived from female donors exhibited early and faster cluster formation, as indicated by higher cluster counts (more clusters/cluster-forming cell) at 2-3 days post-seeding compared to male-donor-derived pseudo-islets. Male-donor-derived pseudo-islets showed overall lower mean cluster counts (fewer clusters/cluster-forming cell), which indicates that each cluster contained more cells. Over longer culture periods (9–10 d), signs of fragmentation emerge, particularly in female-derived pseudo-islets treated with GLP-1. Increased subcluster percentages and a decline in cluster counts indicate this. In male-derived pseudo-islets, fragmentation tends to occur earlier, often by 6–7 days. This is also most pronounced in the + GLP-1 condition. In contrast, GCG supplementation was mostly beneficial for pseudo-islet formation and helped shift toward larger clusters. These findings are consistent with prior co-culture studies, which demonstrated that the presence of GCG- and GLP–1–secreting cells enhances β-cell insulin secretion, proliferation, and resilience to cytotoxic insults (34). However, our study adds the nuance that the supplementation benefit is highly dependent on donor sex and, most importantly, the cellular composition of the islet being formed.

Although hormone supplementation led to notable changes in islet morphology, we did not observe any corresponding differences in hormone secretion from native islets, regardless of sex. This indicates that the structural changes induced by GCG or GLP-1 are not functionally harmful and may in fact reflect a degree of functional resilience in islet hormone output. It is important to consider that islet structure and function are not always tightly linked; alterations such as fragmentation or changes in cluster size may represent adaptive responses to the culture environment rather than early signs of dysfunction. These morphological adaptations could support survival or remodeling without compromising secretory capacity. Additionally, our current assays may not capture subtle changes in function, or the time frame of our culture conditions may be too short to reveal delayed functional effects. Therefore, while morphology was affected, these changes do not appear to impair endocrine function under the conditions tested.

Differences in receptor expression, as visualized in Fig. 5f, were originally hypothesized as the cause of seeing differences in the way cell lines, islets, and pseudo-islets respond. However, although they provide additional insight into the cellular composition and availability of the receptors, there does not seem to be a significant difference between female and male donors. Pseudo-islets exhibited more intense GCGR and GLP-1R staining compared to native islets. This may be due to selection bias introduced during FACS sorting, which enriches for endocrine cells and therefore increases the relative density of receptor-expressing cells in pseudo-islets. It is also possible that improved epitope accessibility in smaller pseudo-islets enhances staining intensity, consistent with their more compact structure and reduced extracellular matrix components.

Finally, when comparing our data on receptor-expressing populations with published datasets, we note that previous reports using mass cytometry (CyTOF) have often described higher proportions of GCGR+ and GLP-1R+ cells (35). However, these differences likely stem from methodological discrepancies, as our study employed FACS-based quantification, which may underrepresent certain populations due to gating thresholds or epitope availability.

Overall, our findings emphasize the importance of selecting the appropriate model system based on the biological process under investigation. Native islets represent the mature, final state of islet architecture and are therefore limited in their utility for studying the effects of hormone supplementation during islet formation. In contrast, pseudo-islets provide a dynamic and controllable model that more accurately captures early stages of reaggregation and cellular adaptation. Our results further underscore the need to consider sex, cellular context, and methodological approach when evaluating islet function and receptor expression.

## Acknowledgments

K.K. and S.P. researched data for cell line experiments. K.K. wrote the manuscript, created the illustrations, and researched and analyzed data for human islet experiments. C.S. analyzed data and reviewed and edited the manuscript. C.D. and P.C. contributed to discussions and helped with FACS data acquisition. Prof. Show-Ling Shyng (OHSU) contributed to discussions and reviewed the manuscript. C.S. is the guarantor for this work.

Human pancreatic islets were provided by the NIDDK-funded Integrated Islet Distribution Program at City of Hope and funded by the IIDP new investigator fund Study #BS562P. Additional human islets were provided by the Alberta Diabetes Institute IsletCore at the University of Alberta in Edmonton (http://www.bcell.org/adi-is-letcore.html) with the assistance of the Human Organ Procurement and Exchange (HOPE) program, Trillium Gift of Life Network (TGLN), and other Canadian organ procurement organizations. FACS was performed with the assistance and facilities of the OHSU Flow Cytometry and Monoclonal Antibody Shared Resource (RRID:SCR_009974). We are grateful for using the Keyence microscope of Prof. David Ellison, Dept. of Medicine, OHSU.

## Notes

https://doi.org/10.5281/zenodo.16757638

https://doi.org/10.5281/zenodo.16757674

### Competing Interest Statement

The authors have declared no competing interest.

## References

1. Tornehave D, Kristensen P, Rømer J, Knudsen LB, Heller RS. Expression of the GLP-1 Receptor in Mouse, Rat, and Human Pancreas. J Histochem Cytochem. 2008 Sep 1;56(9):841–51.

2. Friedrichsen BN, Neubauer N, Lee YC, Gram VK, Blume N, Petersen JS, et al. Stimulation of pancreatic β-cell replication by incretins involves transcriptional induction of cyclin D1 via multiple signalling pathways. Journal of Endocrinology. 2006 Mar 1;188(3):481–92.

3. Kapodistria K, Tsilibary EP, Kotsopoulou E, Moustardas P, Kitsiou P. Liraglutide, a human glucagon-like peptide-1 analogue, stimulates AKT-dependent survival signalling and inhibits pancreatic β-cell apoptosis. Journal of Cellular and Molecular Medicine. 2018;22(6):2970–80.

4. Lee YS, Lee C, Choung JS, Jung HS, Jun HS. Glucagon-Like Peptide 1 Increases b-Cell Regeneration by Promoting a- to b-Cell Transdifferentiation. 2018;67.

5. Thorel F, Népote V, Avril I, Kohno K, Desgraz R, Chera S, et al. Conversion of adult pancreatic alpha-cells to beta-cells after extreme beta-cell loss. Nature. 2010 Apr 22;464(7292):1149–54.

6. Baggio LL, Drucker DJ. Biology of incretins: GLP-1 and GIP. Gastroenterology. 2007 May;132(6):2131–57.

7. Campbell SA, Johnson J, Light PE. Evidence for the existence and potential roles of intra-islet glucagon-like peptide-1. Islets. 13(1–2):32–50.

8. Ho KH, Barmaver SN, Hu R, Yagan M, Ahmed HK, Kaverina I, et al. Pancreatic islet α cells regulate microtubule stability in neighboring β cells to tune insulin secretion and induce functional heterogeneity in individual mouse and human islets [Internet]. bioRxiv; 2024 [cited 2024 Nov 18]. p. 2024.10.21.619544. Available from: https://www.bio-rxiv.org/content/10.1101/2024.10.21.619544v1

9. Bracey KM, Gu G, Kaverina I. Microtubules in Pancreatic β Cells: Convoluted Roadways Toward Precision. Front Cell Dev Biol [Internet]. 2022 Jul 8 [cited 2025 May 18];10. Available from: https://www.frontiersin.org/journals/cell-and-developmental-biology/articles/10.3389/fcell.2022.915206/full

10. Trogden KP, Lee J, Bracey KM, Ho KH, McKinney H, Zhu X, et al. Microtubules regulate pancreatic β-cell heterogeneity via spatiotemporal control of insulin secretion hot spots. Zaidi M, Barton M, Hodson DJ, Bogan JS, editors. eLife. 2021 Nov 16;10:e59912.

11. Tong JCL, Frazer-Morris C, Shilleh AH, Viloria K, de Bray A, Nair AM, et al. Localized GLP1 receptor pre-internalization directs pancreatic alpha cell to beta cell communication. Cell Metab. 2025 Aug 5;37(8):1698–1714.e5.

12. Wieland FC, Sthijns MMJPE, Geuens T, Van Blitterswijk CA, Lapointe VLS. The Role of Alpha Cells in the Self-Assembly of Bioengineered Islets. Tissue Engineering - Part A. 2021 Aug;27(15–16):1055–63.

13. Ast J, Nasteska D, Fine NHF, Nieves DJ, Koszegi Z, Lanoiselée Y, et al. Revealing the tissue-level complexity of endogenous glucagon-like peptide-1 receptor expression and signaling. Nat Commun. 2023 Jan 18;14(1):301.

14. Thiel G, Rössler OG. Glucose Homeostasis and Pancreatic Islet Size Are Regulated by the Transcription Factors Elk-1 and Egr-1 and the Protein Phosphatase Calcineurin. Int J Mol Sci. 2023 Jan 3;24(1):815.

15. Campbell JE, Newgard CB. Mechanisms controlling pancreatic islet cell function in insulin secretion. Nat Rev Mol Cell Biol. 2021 Feb;22(2):142–58.

16. Holter MM, Saikia M, Cummings BP. Alpha-cell paracrine signaling in the regulation of beta-cell insulin secretion. Frontiers in Endocrinology. 2022 Jul 26;13:934775.

17. Bosco D, Armanet M, Morel P, Niclauss N, Sgroi A, Muller YD, et al. Unique Arrangement of ␣- and ␤-Cells in Human Islets of Langerhans.

18. Cabrera O, Berman DM, Kenyon NS, Ricordi C, Berggren PO, Caicedo A. The unique cytoarchitecture of human pancreatic islets has implications for islet cell function. Proceedings of the National Academy of Sciences of the United States of America. 2006 Feb;103(7):2334–9.

19. Lehrstrand J, Davies WIL, Hahn M, Korsgren O, Alanentalo T, Ahlgren U. Illuminating the complete ß-cell mass of the human pancreas-signifying a new view on the islets of Langerhans. Nat Commun. 2024 Apr 18;15(1):3318.

20. Aamodt KI, Powers AC. Signals in the pancreatic islet microenvironment influence β-cell proliferation. Diabetes, Obesity and Metabolism. 2017;19(S1):124–36.

21. Samols E, Weir GC, Bonner-Weir S. Intraislet Insulin-Glucagon-Somatostatin Relationships. In: Lefebvre PJ, editor. Glucagon II [Internet]. Berlin, Heidelberg: Springer Berlin Heidelberg; 1983. p. 133–73. Available from: 10.1007/978-3-642-69019-8_9

22. Richards OC, Raines SM, Attie AD. The role of blood vessels, endothelial cells, and vascular pericytes in insulin secretion and peripheral insulin action. Endocr Rev. 2010 Jun;31(3):343–63.

23. Cleaver O, Dor Y. Vascular instruction of pancreas development. Development. 2012 Aug 15;139(16):2833–43.

24. Lehmann R, Zuellig RA, Kugelmeier P, Baenninger PB, Moritz W, Perren A, et al. Superiority of small islets in human islet transplantation. Diabetes. 2007 Mar;56(3):594–603.

25. Lyon JG, Carr AL, Smith NP, Marfil-Garza B, Spigelman AF, Bautista A, et al. Human research islet cell culture outcomes at the Alberta Diabetes Institute IsletCore. Islets. 2024 Dec 31;16(1):2385510.

26. Abdelli S, Ansite J, Roduit R, Borsello T, Matsumoto I, Sawada T, et al. Intracellular Stress Signaling Pathways Activated During Human Islet Preparation and Following Acute Cytokine Exposure. Diabetes. 2004 Dec 1;53:2815–23.

27. Lyon J, Manning Fox JE, Spigelman AF, Kim R, Smith N, O’Gorman D, et al. Research-Focused Isolation of Human Islets From Donors With and Without Diabetes at the Alberta Diabetes Institute IsletCore. Endocrinology. 2016 Feb;157(2):560–9.

28. Zheng Z, Zong Y, Ma Y, Tian Y, Pang Y, Zhang C, et al. Glucagon-like peptide-1 receptor: mechanisms and advances in therapy. Sig Transduct Target Ther. 2024 Sep 18;9(1):1–29.

29. Efrat S. Ex-vivo Expansion of Adult Human Pancreatic Beta-Cells. Rev Diabet Stud. 2008;5(2):116–22.

30. Dorrell C, Schug J, Canaday PS, Russ HA, Tarlow BD, Grompe MT, et al. Human islets contain four distinct sub-types of β cells. Nature Communications. 2016;7:1–9.

31. Müller TD, Finan B, Bloom SR, D’Alessio D, Drucker DJ, Flatt PR, et al. Glucagon-like peptide 1 (GLP-1). Mol Metab. 2019 Sep 30;30:72–130.

32. Nadal A, Alonso-Magdalena P, Soriano S, Ropero AB, Quesada I. The role of oestrogens in the adaptation of islets to insulin resistance. J Physiol. 2009 Nov 1;587(Pt 21):5031–7.

33. Soriano S, Alonso-Magdalena P, García-Arévalo M, Novials A, Muhammed SJ, Salehi A, et al. Rapid insulinotropic action of low doses of bisphenol-A on mouse and human islets of Langerhans: role of estrogen receptor β. PLoS One. 2012;7(2):e31109.

34. Green AD, Vasu S, Moffett RC, Flatt PR. Co-culture of clonal beta cells with GLP-1 and glucagon-secreting cell line impacts on beta cell insulin secretion, proliferation and susceptibility to cytotoxins | Elsevier Enhanced Reader. Biochimie. 2016 Mar 22;125:119–25.

35. Wang YJ, Golson ML, Schug J, Traum D, Liu C, Vivek K, et al. Single-Cell Mass Cytometry Analysis of the Human Endocrine Pancreas. Cell Metab. 2016 Oct 11;24(4):616–26.

