## Supplemental Data for "Glucagon and GLP-1 Accelerate Pseudo-Islet Assembly and Unmask Sex-Specific Islet Fragmentation Dynamics"

### Supplements

### Instruments

Instrumentation and assay analysis were performed using standardized and optimized protocols across multiple platforms. Confocal imaging was conducted on an Olympus Fluoview 1200 dual-scanner confocal microscope using a 60x oil objective. Fluorescence detection was achieved using the following laser lines and power settings: 405 nm at 2.0 % laser power (LP) for the blue channel, 488 nm at 2.0 % LP for the green channel, 559 nm at 5.0 % LP for the yellow channel, and 635 nm at 5.0 % LP for the red channel. EPI-fluorescence imaging was carried out on a BZ-X800 microscope (Keyence) in brightfield mode, equipped with a CFI Plan Apo 4x objective. Imaging was performed in plastic-bottom 6-well plates. This combination has a high-resolution pixel ratio of 1.88721 µm/pixel.

Flow cytometric cell sorting (FACS) was primarily conducted on a Cytopeia inFluxV-GS (Becton-Dickinson, BD) using propidium iodide (PI) for dead cell exclusion, and fluorescently labeled antibodies conjugated to Alexa Fluor 488 (A488), Phycoerythrin (PE), and Allophycocyanin (APC). PI, A488, and PE were excited using a 488 nm, 100 mW laser, while APC was excited using a 638 nm, 100 mW laser. When the inFluxV-GS was unavailable, sorting was performed on a BD Symphony S6, using the same staining panel. However, APC was excited with a 637 nm, 140 mW laser, and PE was excited with a 561 nm, 100 mW laser. Enzyme-linked immunosorbent assays (ELISAs) were performed using Promega LUMIT kits for insulin (CS3037A05) and glucagon (W8020). All assays were conducted according to manufacturer instructions, with modifications to reaction volumes for 384-well plate formats based on optimization guidance provided by Dr. M. Cappozzi (personal communication).

**Supplemental Table 1: Donor clinical parameters.** All donors are identified by donor ID, 8 digits for IIDP-derived donors and R-XXX (3 digits) for ADI-derived donors. Donors are grouped by sex (with 17 of each gender). Additionally, donor age, BMI, and % A1C are listed. Percent β-cells, α-cells, and δ-cells represent the cell percentage yield post-FACS. Donors listed as N/A were not sorted, and only native islets were used in experiments.

| Donor ID | Gender | Type | Age | BMI | % A1C | % β-cells | % α-cells | % δ-cells |
|---|---|---|---|---|---|---|---|---|
| 36823227 | Female | No Disease (ND) | 41 | 38.2 | 5.2 | 17.34 | 1.48 | 3.26 |
| 37350251 | Female | ND | 53 | 29.9 | 5.2 | N/A* | N/A | N/A |
| $R525_{ADI}$ | Female | ND | 51 | 47.8 | 6.5 | N/A | N/A | N/A |
| 40709610 | Female | ND | 55 | 28.6 | 5.7 | N/A | N/A | N/A |
| 39980165 | Female | ND | 60 | 29.6 | 6.2 | 23.3 | 5.8 | 1.93 |
| 40992344 | Female | ND | 42 | 33.3 | 5.5 | N/A | N/A | N/A |
| 44337332 | Female | ND | 31 | 16.3 | 6 | 8.66 | 8.47 | 2.7 |
| 43583143 | Female | ND | 19 | 22.3 | 6 | 6.98 | 0.95 | 8.33 |
| 43463703 | Female | ND | 54 | 28.7 | 5.7 | | | |
| $R551_{ADI}$ | Female | ND | 57 | 26 | 4.3 | 5.1 | 0.21 | 10.9 |
| $R546_{ADI}$ | Female | ND | 42 | 29.7 | 5.9 | 18.14 | 8.87 | 3.5 |
| $R537_{ADI}$ | Female | ND | 18 | 37.3 | 5.5 | | | |
| $R532_{ADI}$ | Female | ND | 28 | 24.3 | 5.5 | 16.1 | 0.2 | 1.2 |
| $R510_{ADI}$ | Female | ND | 53 | 21.3 | 5.9 | 7.29 | 0.42 | 0.77 |

| | | | | | | | | |
|---|---|---|---|---|---|---|---|---|
| 47290391 | Female | ND | 54 | 30.9 | 4.7 | N/A | N/A | N/A |
| R552$_{ADI}$ | Female | ND | 69 | 19.5 | 6.5 | N/A | N/A | N/A |
| R575$_{ADI}$ | Female | ND | 59 | 43.9 | 6.1 | N/A | N/A | N/A |
| 39708909 | Male | ND | 21 | 38.1 | 5 | 33.6 | 22 | 1.76 |
| 38750927 | Male | ND | 34 | 24.5 | 5.4 | 12.02 | 27.1 | 3.5 |
| 36510137 | Male | ND | 63 | 32.1 | 6.1 | N/A | N/A | N/A |
| R549$_{ADI}$ | Male | ND | 54 | 26.5 | 5.1 | 8.8 | 0.3 | 0.13 |
| 28501433 | Male | ND | 48 | 32.3 | 5.7 | 28 | 24 | 2 |
| 43079887 | Male | ND | 43 | 24.1 | 4.7 | 12 | 18 | 0.24 |
| 42355911 | Male | ND | 64 | 29.5 | 5.7 | 28.7 | 9.2 | 3.8 |
| 40380406 | Male | ND | 47 | 34.2 | 5.5 | 10.4 | 6.6 | 5.8 |
| R495$_{ADI}$ | Male | ND | 50 | 33.8 | 5.8 | 7.58 | 0.43 | 2.93 |
| 43897604 | Male | ND | 41 | 27.8 | 5.3 | 0.57 | 0.18 | 0.22 |
| 42160709 | Male | ND | 49 | 26 | 4.8 | 20.13 | 6.47 | 5.85 |
| 42008301 | Male | ND | 36 | 30 | 5 | 12.6 | 6 | 4.2 |
| 41299360 | Male | ND | 20 | 24 | 5.1 | 15.5 | 6.3 | 5.7 |
| 39708909 | Male | ND | 21 | 38.1 | 5 | 12.7 | 12.1 | 4.9 |
| 37638596 | Male | ND | 34 | 27 | 5.5 | 20.1 | 3.6 | 2.1 |
| R501$_{ADI}$ | Male | ND | 28 | 23.9 | 4.7 | 14.81 | 2.21 | 0.15 |
| 46795533 | Male | ND | 58 | 39.2 | 5.9 | N/A | N/A | N/A |

* not sorted

**Supplemental Table 2: Summary of pseudo-islet experimental conditions and interpretations.** This table summarizes the interpretation of pseudo-islet data across all experimental conditions, including variations in cell composition, days post-seeding, and media supplementation. Due to the increased complexity compared to native islets, this comprehensive overview is provided in the supplement to conserve space in the main text, which includes one representative example.

| Donor gender | Cell combination | Media supplementation | Time post-seeding | Mean cluster count (dynamic growth) | % subclusters | Interpretation |
|---|---|---|---|---|---|---|
| Female | α+β+δ:MS1 | None | 2-3 days | 0.0157 | 100 | Baseline |
| Male | | | | 0.0108 | 100 | |
| Female | | + GCG | | 0.0250 | 96.1 | |
| Male | | | | 0.0113 | 71.9 | |
| Female | | + GLP-1 | | - | - | |
| Male | | | | 0.0103 | 89.2 | |
| Female | | None | 6-7 days | 0.0107 | 100 | - |
| Male | | | | 0.0095 | 100 | - |
| Female | | + GCG | | 0.0122 | 42.6 | Cluster ⇓<br>% subcluster ⇓<br>= fusion into larger clusters |
| Male | | | | 0.0122 | 172.9 | Cluster ⇑<br>% subcluster ⇑<br>= increased formation of small aggregates |
| Female | | + GLP-1 | | - | - | - |
| Male | | | | 0.0089 | 114.1 | Cluster ⇓<br>% subcluster ⇑ |

| | | | | | | |
|---|---|---|---|---|---|---|
| | | | | | | = fragmentation |
| Female | | None | 9-10 days | - | - | - |
| Male | | | | 0.0084 | 100 | - |
| Female | | + GCG | | - | - | - |
| Male | | | | 0.0114 | 108.3 | Cluster ⇓ % subcluster ⇓ = fusion into larger clusters |
| Female | | + GLP-1 | | - | - | - |
| Male | | | | 0.0052 | 85.9 | Cluster ⇓ % subcluster ⇓ = fusion into larger clusters |
| Female | β+δ:MS1 | None | 2-3 days | 0.0181 | 100 | Baseline |
| Male | | | | 0.0165 | 100 | |
| Female | | + GCG | | 0.0114 | 53.2 | |
| Male | | | | 0.0188 | 74.6 | |
| Female | | None | 6-7 days | 0.0362 | 100 | - |
| Male | | | | 0.0051 | 100 | - |
| Female | | + GCG | | 0.0088 | 22.2 | Cluster ⇓ % subcluster ⇓ = fusion into larger clusters |
| Male | | | | 0.0074 | 68 | Cluster ⇓ % subcluster ⇓ = fusion into larger clusters |
| Female | | None | 9-10 days | 0.0615 | 100 | - |
| Male | | | | 0.0049 | 100 | - |
| Female | | + GCG | | 0.0450 | 54.3 | Cluster ⇑ % subcluster ⇑ = increased formation of small aggregates |
| Male | | | | 0.0117 | 38.7 | Cluster ⇑ % subcluster ⇓ = shift toward larger clusters |
| Female | α+β:MS1 | None | 2-3 days | 0.0029 | 100 | Baseline |
| Male | | | | 0.0147 | 100 | |
| Female | | + GCG | | 0.0148 | 334 | |
| Male | | | | 0.0086 | 33.1 | |
| Female | | + GLP-1 | | 0.0023 | 370.5 | |
| Male | | | | 0.0107 | 60.5 | |
| Female | | None | 6-7 days | 0.0044 | 100 | - |
| Male | | | | 0.0050 | 100 | - |
| Female | | + GCG | | 0.0031 | 48 | Cluster ⇓ % subcluster ⇓ = fusion into larger clusters |

| | | | | | | |
|---|---|---|---|---|---|---|
| Male | | | | 0.0049 | 65 | Cluster ⇓<br>% subcluster ⇑<br>= fragmentation |
| Female | | + GLP-1 | | 0.0203 | 173.8 | Cluster ⇑<br>% subcluster ⇓<br>= shift toward larger clusters |
| Male | | | | 0.0087 | 193.2 | Cluster ⇓<br>% subcluster ⇑<br>= fragmentation |
| Female | | None | 9-10 days | 0.0035 | 100 | - |
| Male | | | | - | - | - |
| Female | | + GCG | | - | - | - |
| Male | | | | - | - | - |
| Female | | + GLP-1 | | 0.0016 | 686.6 | Cluster ⇓<br>% subcluster ⇑<br>= fragmentation |
| Male | | | | 0.0101 | - | - |

Wang et al. (1) have used single-cell mass spectrometry (cyTOF) to quantify the % of endocrine cell populations within human islets. This was done across 17 healthy donors (7 female and 10 male) and 3 T2D donors (1 female, 2 male). The data generated in their study serves as the literature reference for our results and is summarized in Supplemental Figure 1a.

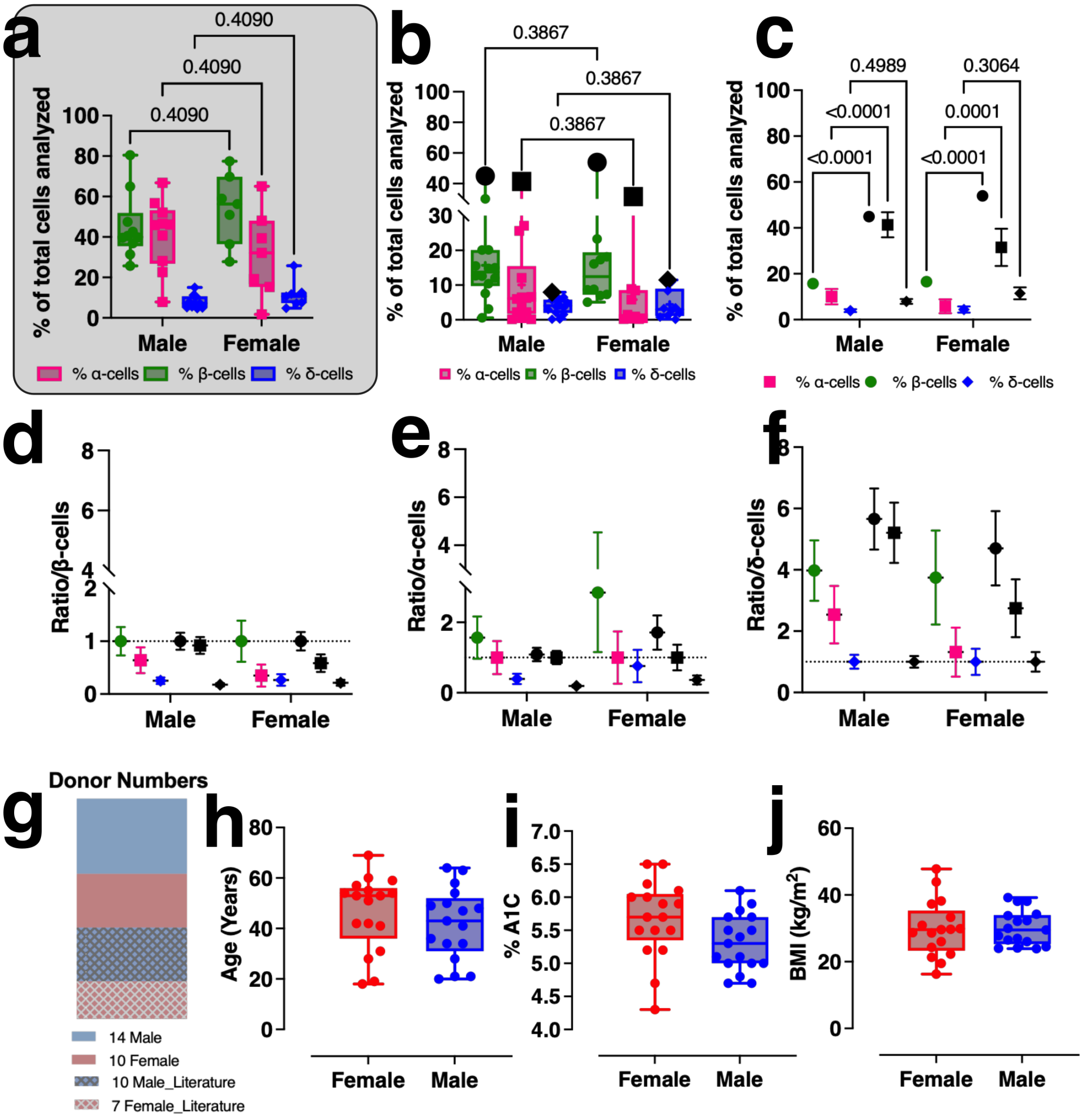


**Supplemental Figure 1: Comparison of this study's FACS results with available literature.** a) adapted from Wang *et al.* visualizing % of α-cells (pink), β-cells (green) and δ-cells (blue) of total cells analyzed by single cell mass cytometry (cyTOF) for both male and female healthy donors. b) Percentage of α-cells (pink), β-cells (green), and δ-cells (blue) of total cells analyzed by FACS in the present study. The mean literature value for each cell type is depicted in black. c) Percentage of α-cells (pink), β-cells (green), and δ-cells (blue) for all cells analyzed by FACS vs. cyTOF (literature). Literature values follow the same symbol association but are labeled black. d-f) Ratio of cells expressed as /β-cells (d), /α-cells (e), and /δ-cells (f) for healthy male and female donors evaluated in this study and in comparison to literature results (following the same shape association but colored black). g) Pie chart of the number of healthy male (blue) and

**Fluorescence-activated cell sorting (FACS) procedure for pseudo-islet generation**

Following the pre-established protocol from the Grompe lab (2), dissociated human pancreatic islets were sorted into pure α-, β-, and δ-cell populations. Islet cells were first separated from debris based on their position along the forward scatter axis (Supplemental Fig. 2a). Non-single cells were excluded using the Trigger Pulse Width parameter (Supplemental Fig. 2b). Dead cells were removed by propidium iodide staining (Supplemental Fig. 2c). Antibody staining with HIC1-2B4, HIC1-8G12, and CD9 was used to distinguish α-cell/δ-cell and β-cell/δ-cell pre-populations, which were subsequently further refined (Supplemental Fig. 2d–f).

For each sort, performance was evaluated by using cells that were not associated with any of the desired populations. For this, a sample of those unassociated cells was collected and re-sorted with the same settings. In case of a clean and pure sort, a negligible percentage (> 0.5 %) of the sorted cells appear in the α-, β-, or δ-cell gates (see Supplemental Fig. 2g and h). Additionally, by specifically gating for propidium iodide-positive (PI+) cells,

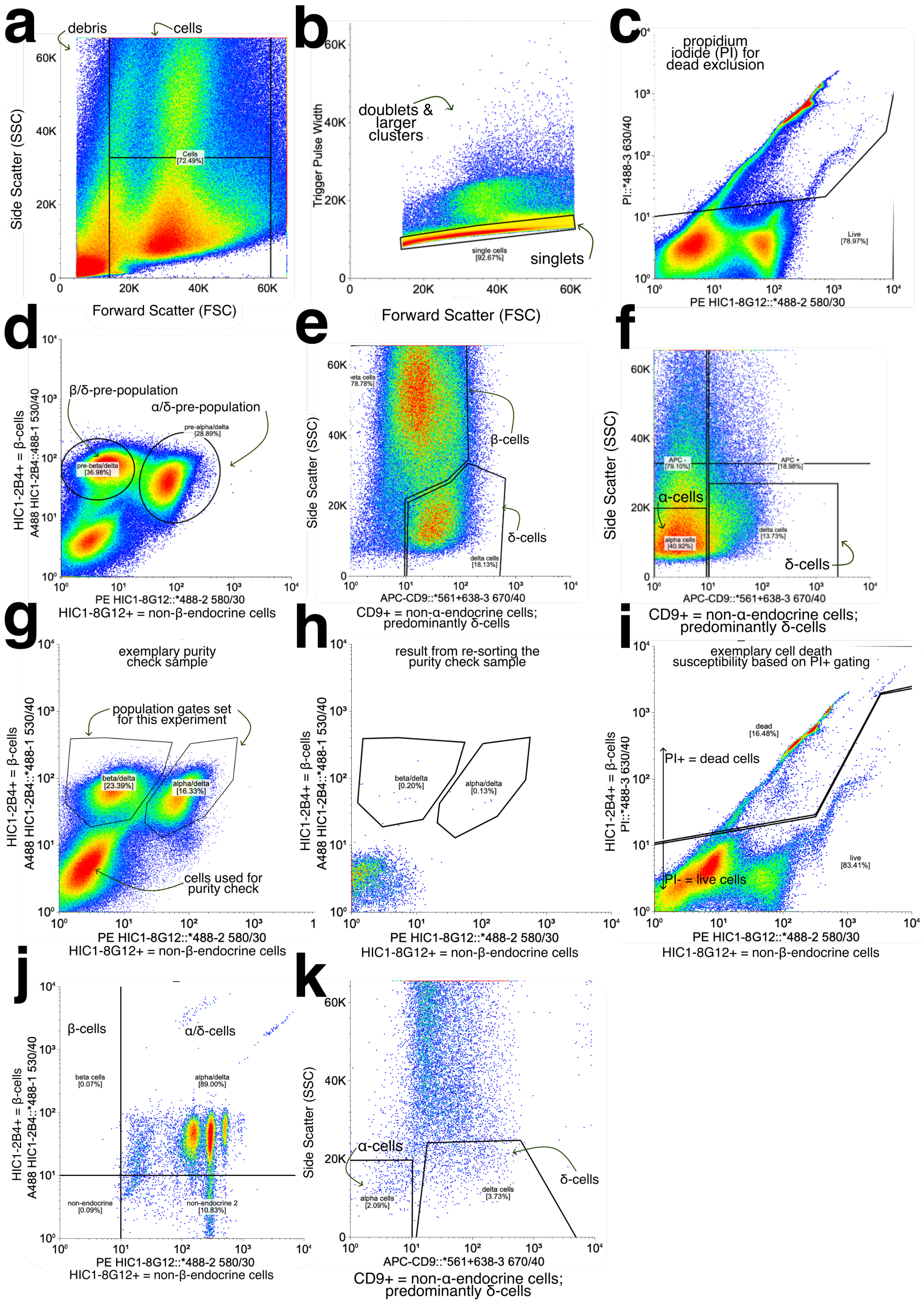


**Supplemental Figure 2: Cell sorting (FACS) workflow for isolating endocrine cell populations from human donor islets.** Human native islets are enzymatically dissociated into single-cell suspensions and stained with a combination of endocrine-specific antibodies: HIC1-2B4, which labels β-cells; HIC1-8G12, which labels non-β endocrine cells; and CD9, which labels δ-cells and a minor subset of β-cells, and is used to exclude α-cells. This antibody panel allows for the identification and separation of β-cells, α-cells, and δ-cells. a) Debris is excluded based on forward scatter (FSC) and side scatter (SSC) parameters. b) Single cells are gated by excluding doublets and aggregates. c) Propidium iodide (PI) is used to exclude dead cells; only PI-negative (viable) cells are analyzed. d) Cells are first separated based on HIC1-2B4 (y-axis) and HIC1-8G12 (x-axis) signal to identify prepopulations of β-/δ-cells and α-/δ-cells. e, f) Within these prepopulations, CD9 expression and SSC are used to further resolve β-cells from δ-cells (e) and α-cells from δ-cells (f). Before sorting, 10 µm reference beads are used to identify the cell population window. Non-endocrine or undefined cells are collected separately for post-sort purity assessment. All FACS data were analyzed using Floreada.io.

we were able to identify the percentage of cells excluded and the especially susceptible populations. By doing so, we have identified that ~ 20 % of cells are excluded as dead. From those 20 %, 0.07 % are β-cells, 2 % are α-cells, and 3.7 % are δ-cells; recalculated for all cells analyzed, this translates to 0.007 % β-cells, 0.16 % α-cells, and 0.31 % δ-cells (see Supplemental Fig. 2i-k).

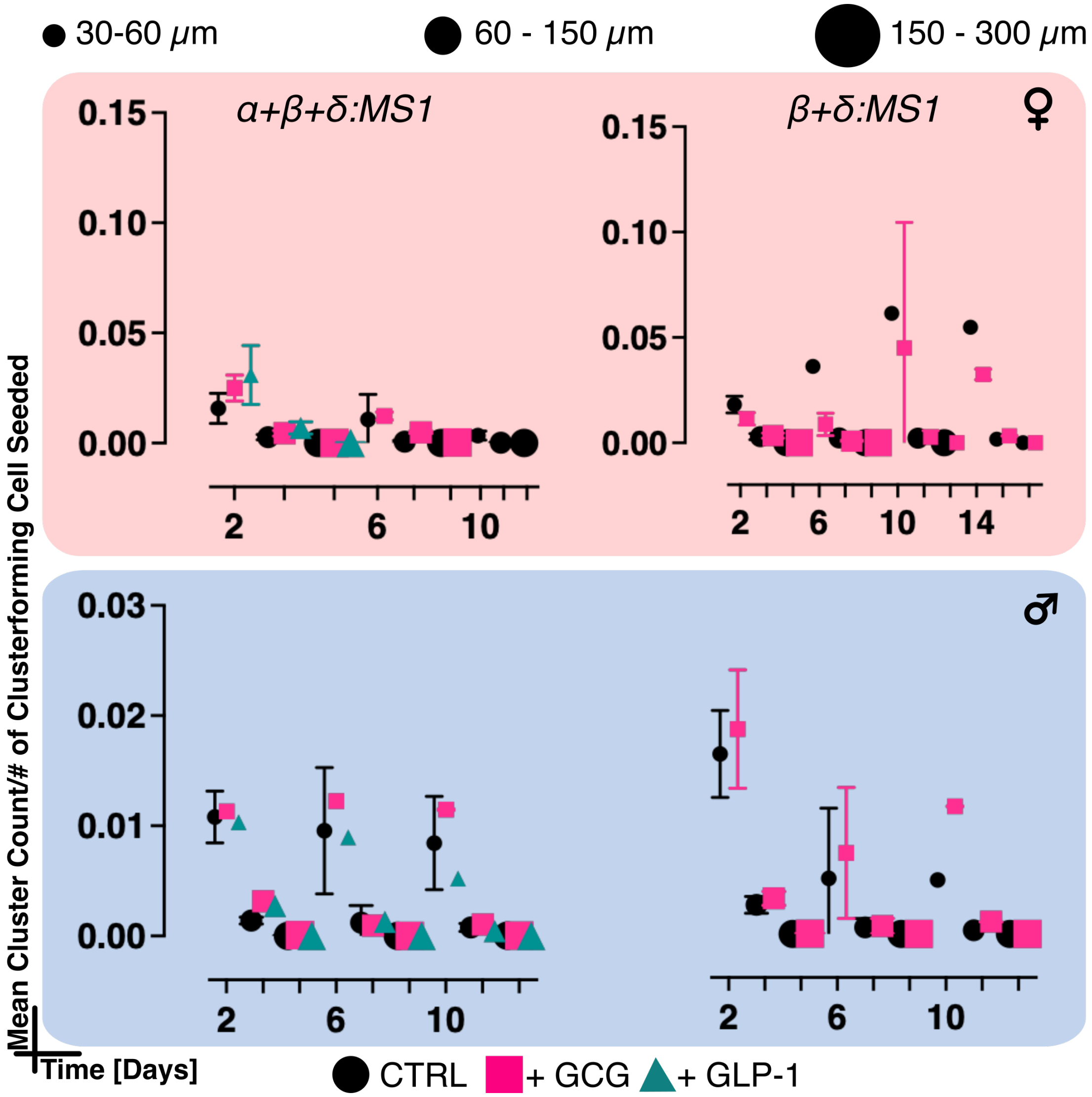


**Supplemental Figure 3: Pseudo-islet cluster counts monitored over size and time.** Mean pseudo-islet cluster counts normalized to the # of cluster-forming cells seeded in each well were monitored for their time in culture (measured in days). For each day, three different sizing bins were measured: 30-60 μm cluster diameter (small symbols), 60-150 μm diameter (medium symbols), and 150-300 μm diameter (large symbols). Both female (red) and male (blue) pseudo-islet clusters were monitored for cell combinations of α+β+δ:MS1 and β+δ:MS1.

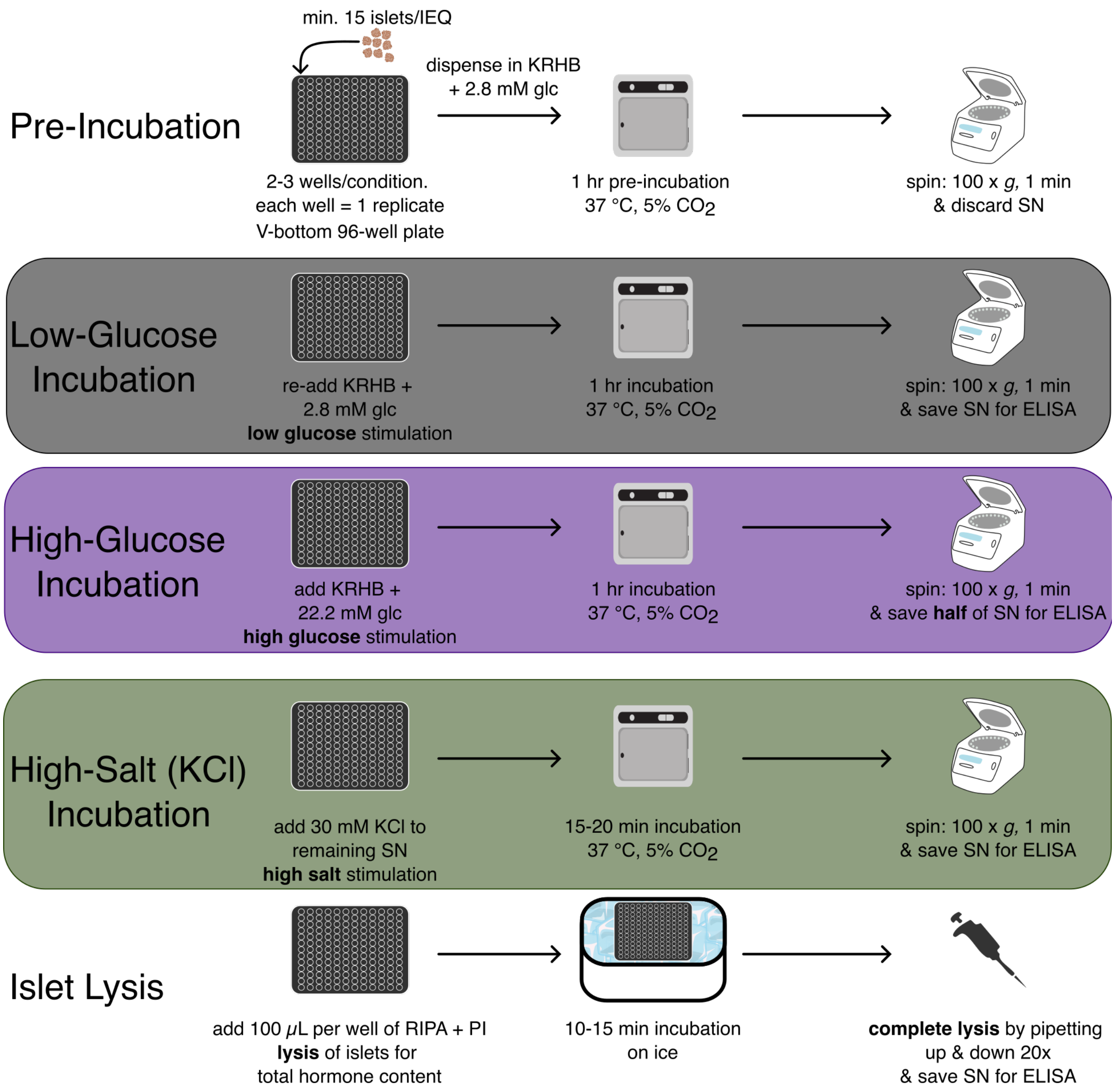


**Supplemental Figure 4: Overview of the glucose-stimulated hormone secretion (GSHS) assay workflow.** To evaluate native and pseudo-islet functionality, a minimum of 15 islet equivalents (IEQ) per sample were manually picked and transferred to a V-bottom 96-well plate. The conical well shape facilitated gentle handling by allowing islets to settle at the bottom, minimizing loss during wash and buffer exchange steps. Each condition was assessed in 2–3 technical replicates. Following transfer, islets were pre-incubated for 1 hour in Krebs-Ringer HEPES buffer (KRHB) containing 2.8 mM glucose to equilibrate to the assay environment. After pre-incubation, plates were centrifuged (1 min, 100 × g), and the supernatant (SN) discarded. The assay was then carried out in three sequential stimulation steps: 1) Low-glucose stimulation (black panel): Islets were incubated in KRHB + 2.8 mM glucose for 1 hour at 37 °C and 5% $CO_2$. After centrifugation, the SN was collected for ELISA. 2) High-glucose stimulation (purple panel): Islets were then incubated in KRHB + 22.2 mM glucose under the same conditions. After 1 hour, half the SN was collected for ELISA. 3) High-potassium depolarization (green panel): KCl was added to the remaining SN to a final concentration of 30 mM. Islets were incubated for an additional 15–20 minutes at 37 °C, then spun down, and the SN collected. To assess total hormone content, islets were lysed in RIPA buffer supplemented with protease inhibitor cocktail (PI) on ice for 10–15 minutes. Lysis was completed by pipetting up and down 20 times. The resulting lysate was saved for total hormone content ELISA analysis. All supernatant and lysate samples were stored at 4 °C and processed within 7 days using PROMEGA Lumit ELISA kits for insulin and glucagon.

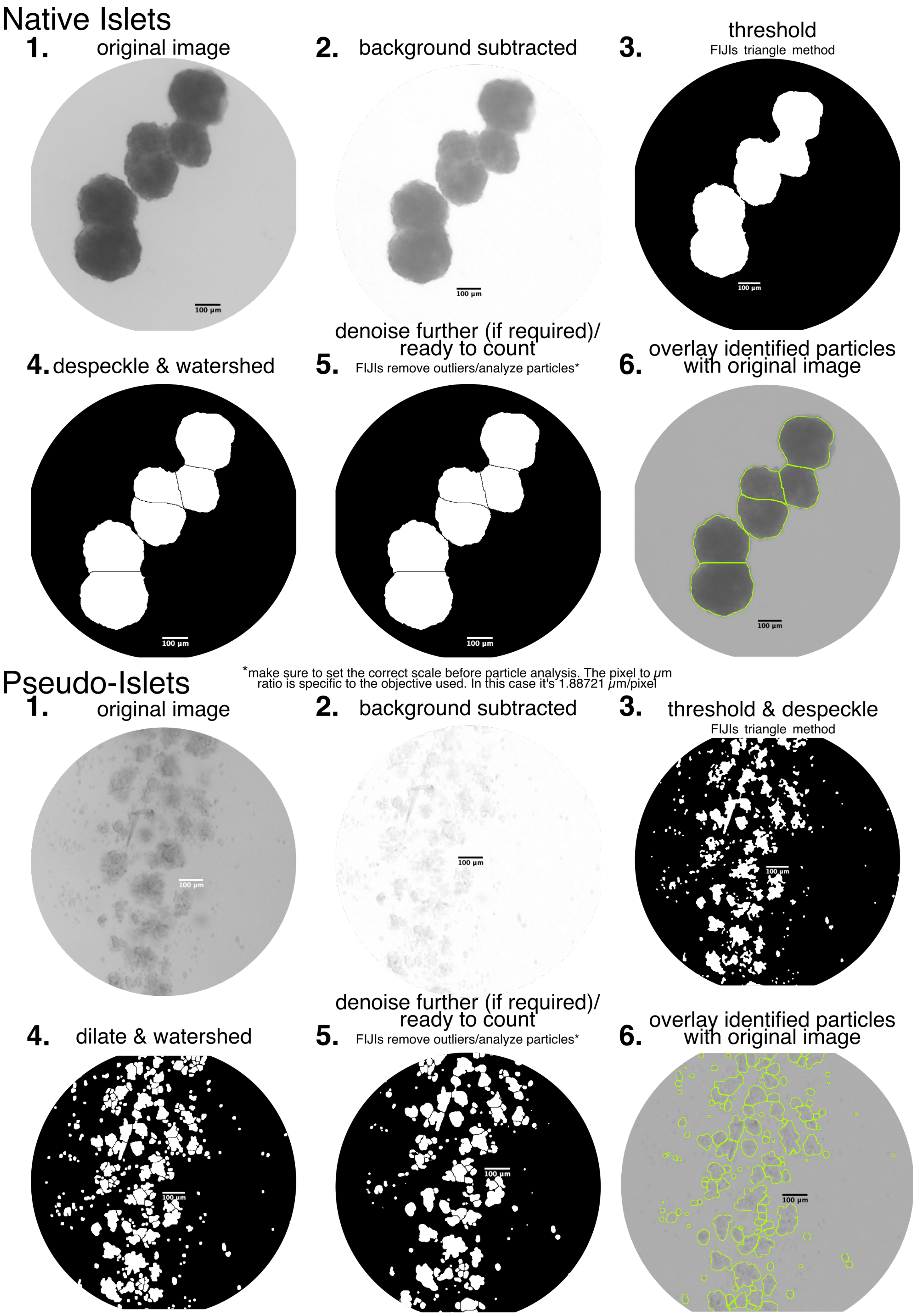


**Supplemental Figure 5: Step-by-step image analysis pipeline in FIJI to quantify native and pseudo-islets.** Native and pseudo-islets were cultured in ultra-low attachment 6-well plates and imaged using a Keyence EPI fluorescence microscope. 1) Original image: Brightfield images capturing the entire well. 2) Background subtraction: FIJIs rolling ball algorithm was applied (radius = 100 px for native islets; 20 px for pseudo-islets) to enhance contrast. 3) Thresholding and despeckling: The Triangle method was used for thresholding, followed by despeckling to reduce background noise. 4) Preprocessing for segmentation: "watershed" was used to separate adjacent native islets. For pseudo-islets, a "dilate" step preceded "watershed" to close internal gaps. 5) Denoising and particle analysis: Outliers were removed, and particle analysis was performed after image scale calibration to count and measure islet-like structures. 6) Validation via overlay: Detected particles were overlaid on the original image to confirm accurate segmentation. Zoomed-in views are shown with a 100 µm scale bar for reference.

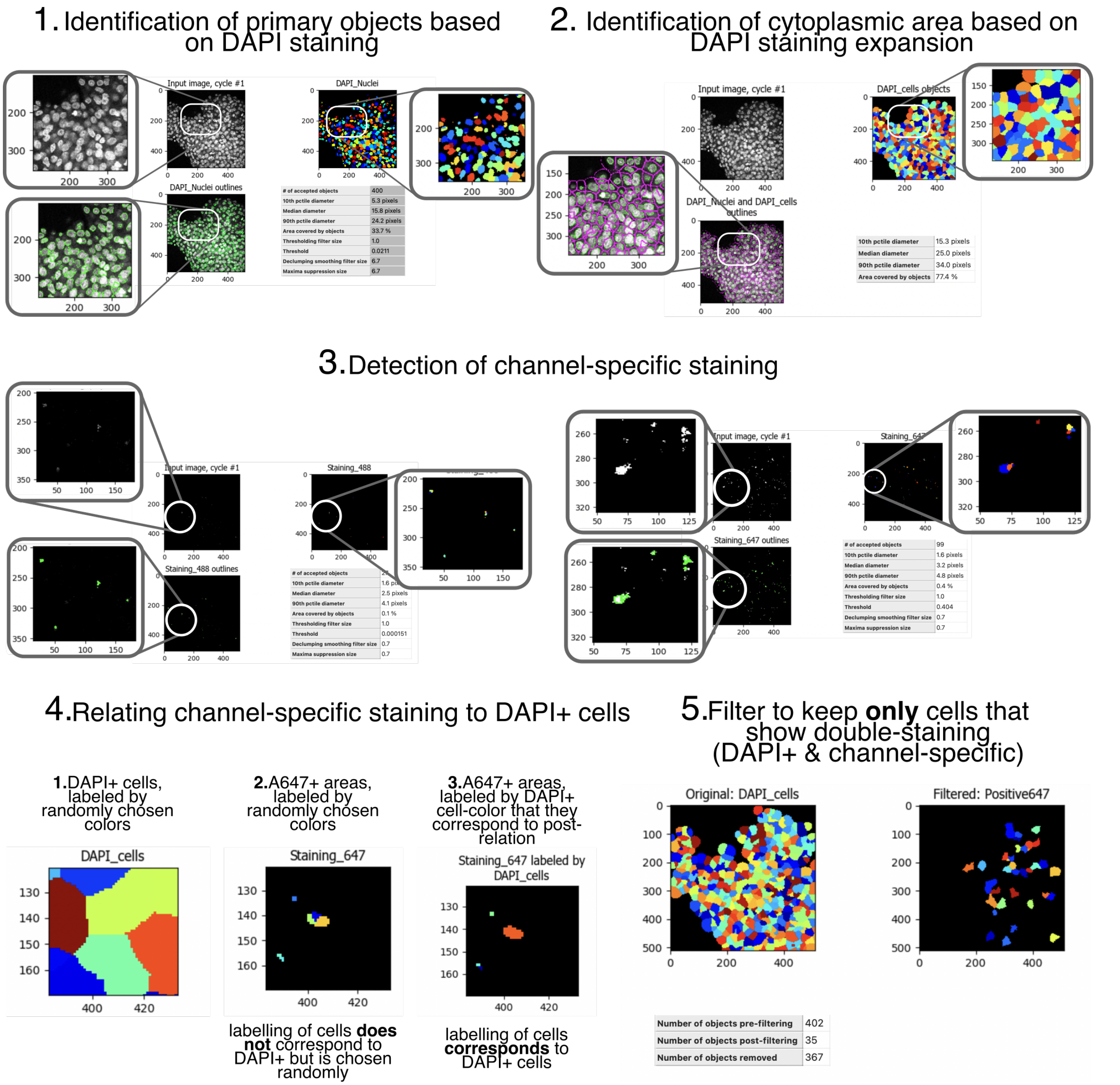


**Supplemental Figure 6: Overview of the CellProfiler pipeline used to quantify GCGR and GLP-1R expressing cells in native and pseudo-islets.** To assess GCGR and GLP-1R expression in human donor-derived native and pseudo-islets, islets were PFA-fixed and immunostained for insulin (559 nm), glucagon receptor (GCGR, 488 nm), and GLP-1 receptor (GLP-1R, 647 nm). Nuclei were stained with DAPI (405 nm). Confocal z-stack images were acquired using a 60× oil objective to capture the 3D structure of each islet. Pre-processing: Images were pre-processed in FIJI. This included channel splitting and brightness/contrast adjustment to match the DAPI channel. For each islet, every 2nd or 3rd plane from the z-stack (depending on islet size) was selected. Grayscale images were saved for each z-plane and for each color combination: 405+488+559, 405+488+647, and 405+559+647. CellProfiler analysis steps: 1) Primary object identification: Nuclei were identified using the DAPI (405 nm) channel staining. 2) Cytoplasmic expansion: A cytoplasmic area was approximated by expanding each nucleus by 15 pixels. 3) Channel-Specific Signal detection: Receptor-specific signal (e.g., 488 nm for GCGR, 559 nm for insulin, and 647 nm for GLP-1R) was detected in the corresponding channels. Examples are shown for 488 nm and 647 nm. 4) Relating objects: Staining-positive areas were mapped to DAPI+ nuclei to assign expression to individual cells. 5) Filtering: Only cells with both a DAPI+ nucleus and detectable receptor signal were retained for analysis; all unmatched objects were excluded. Each color combination was analyzed using a dedicated CellProfiler pipeline. Cell detection was visually verified before steps 3–5 to ensure accurate segmentation and signal attribution.

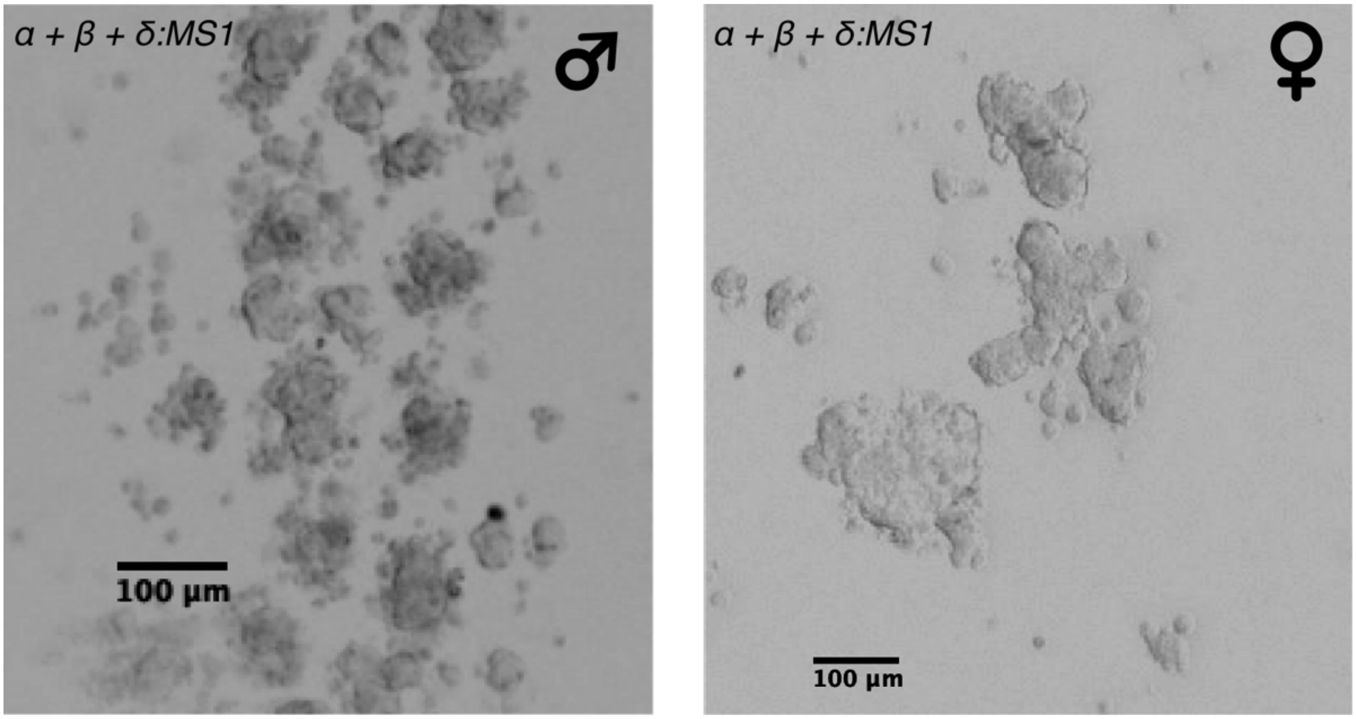


**Supplemental Figure 7: Pseudo-islet brightfield images.** Using a Keyence EPI-fluorescence microscope in brightfield mode, pseudo-islets were imaged every 2-3d to monitor their growth. Left: male-donor-derived pseudo-islets of α+β+δ:MS1 composition at 2-3 d post-seeding. Right: female-donor-derived pseudo-islets of α+β+δ:MS1 composition at 2-3 d post-seeding. Scale bar shown in black corresponds to 100 µm

### References

1. Wang YJ, Golson ML, Schug J, Traum D, Liu C, Vivek K, et al. Single-Cell Mass Cytometry Analysis of the Human Endocrine Pancreas. Cell Metab. 2016 Oct 11;24(4):616–26.

2. Dorrell C, Schug J, Canaday PS, Russ HA, Tarlow BD, Grompe MT, et al. Human islets contain four distinct subtypes of β cells. Nature Communications. 2016;7:1–9.
